# Cognitive function linked to temporal occupancy of Brain-Ventricle (BraVe) modes

**DOI:** 10.1101/2025.01.04.631289

**Authors:** Fulvia Francesca Campo, Vânia Miguel, Elvira Brattico, Salvatore Nigro, Benedetta Tafuri, Giancarlo Logroscino, Joana Cabral, the Alzheimer’s Disease Neuroimaging Initiative (ADNI)

## Abstract

The temporal coordination between fMRI signals in gray matter and ventricular spaces represents an emerging yet poorly understood dimension of brain physiology with potential relevance for neuroprotection. Here, we analyzed 2177 resting-state fMRI scans from 880 participants spanning the cognitive spectrum from normal aging to dementia, using a unified phase-coupling approach that incorporates fluctuations in both brain tissue and ventricular space. We identify distinct modes of Brain-Ventricle (BraVe) coupling occurring over time, and compare their temporal occupancy across cognitive status groups, neurodegenerative biomarkers and cognitive scores.

In the most prevalent BraVe mode, signals across the entire brain evolve in anti-phase with ventricular signals, occurring less frequently with cognitive decline (p < 10^-7^, g > 0.3). In the remaining BraVe modes, specific cortical regions temporarily align with ventricular fluctuations, shaping canonical resting-state networks at the cortical level. Two BraVe modes overlapping with the Default Mode Network and the Frontoparietal Network, occur less frequently in dementia (p < 10^-8^, g > 0.35), with the latter correlating significantly with FDG-PET glucose metabolism (r = 0.20, p = 3.65×10^-5^). Conversely, a mode overlapping with the Limbic Network increases in occupancy with cognitive decline and associates with APOE ε4 status (r = 0.11, p = 1.30×10^-6^).

BraVe mode occupancy correlates more strongly and broadly with cognitive function than with disease-specific molecular biomarkers, with between 16 and 27 cognitive and functional variables per mode surviving strict Bonferroni correction, spanning global cognition, episodic memory, executive function, and everyday functioning. These findings reveal that the transient phase shifts underlying resting-state network dynamics are not arbitrary: when cortical subnetworks shift out of phase from the global brain signal, they align with ventricular signals, suggesting that resting-state functional connectivity and brain-ventricle coupling may be two perspectives on the same underlying physical process, one that is directly linked with cognitive function.

## 1. Introduction

Healthy cognitive function emerges from coordinated activity across distributed brain networks, a fragile architecture vulnerable to aging and disease. When this coordination breaks down, cognitive abilities decline progressively, in many cases evolving into dementia, a condition affecting nearly 4% of people aged 65 and older and placing an enormous burden on patients, families, and healthcare systems worldwide. Despite decades of research, no disease-modifying treatments have proven broadly effective, underscoring the need to revisit the fundamental mechanisms that sustain healthy brain function and render it vulnerable to deterioration.

Traditionally, cognitive decline has been approached from a neurodegenerative perspective, assuming that neuronal death and synaptic loss directly cause cognitive impairment. However, this relationship is more complex than initially thought: some individuals maintain cognitive function despite significant gray matter atrophy and neuropathological burden^1–3^, suggesting that additional mechanisms beyond neuronal integrity contribute to cognitive preservation or vulnerability. This suggests that additional mechanisms beyond neuronal integrity contribute to cognitive preservation or vulnerability. Among them, the dynamics of brain fluids and their interaction with neural tissue, including glymphatic clearance pathways, have emerged as a growing focus of investigation^4,5^.

Studies have demonstrated that cerebrospinal fluid (CSF) inflow into the fourth ventricle negatively correlates with the global BOLD signal in gray matter, a phenomenon termed gBOLD-CSF coupling ^6,7^. This coupling is particularly strong during deep sleep, a finding that has transformed our understanding of the role of sleep in brain clearance^6,8^. Despite being weaker, it is also detected during wakefulness with resting-state fMRI, where it has been shown to decrease with cognitive decline, such as in Alzheimer’s Disease (AD)^9^ and Parkinson’s Disease^10^, challenging the conventional neurocentric paradigm. Critically, current understanding remains limited to this single mode of brain-ventricle coupling. In contrast, the existence and potential significance of more complex brain-ventricle coupling modes for cognitive function remain entirely unexplored.

Both brain clearance mechanisms and resting-state fMRI fluctuations operate in the same ultra-slow frequency range (below 0.1 Hz), with spontaneous activity waves shown to propagate along cortical hierarchical gradients^11,12^. Yet the relationship between these dynamics and ventricular signals remains poorly understood, motivating the exploration of coupling patterns beyond the established global-to-ventricles relationship^13–15^.

Most resting-state fMRI research on cognitive decline has focused exclusively on signal fluctuations in cortical and subcortical gray matter, identifying consistent modes of functional connectivity and co-activation patterns altered across the spectrum of cognitive impairment, from transgenic mouse models of amyloidosis to human cohorts spanning normal cognition, mild cognitive impairment (MCI) and dementia^16–20^. While altered DMN connectivity has been recognised as a candidate imaging biomarker for dementia^21,22^, it currently lacks clinical validation and has been characterised primarily through group-wise comparisons in relatively small single-centre samples, with the optimal analytical approach remaining an open question. The Global Mode, in which signals from all brain areas slowly co-vary, has been found to be most distinctive of cognitive performance^23^, yet other configurations in which a subset of brain regions temporarily synchronize have also been identified. In MCI and dementia, decreased functional connectivity within the Default Mode Network (DMN) is particularly well documented^24–30^, with other networks including the Salience Network^28,31,32^, Central Executive Network (CEN)^33–36^ and subcortical structures^37–40^ also showing alterations with cognitive decline. A fundamental question nonetheless remains unanswered: what coordinates these transient episodes of phase coherence, and why do signals in spatially distributed regions shift out of phase from the rest of the brain in the first place?

Most neuroimaging preprocessing approaches have dismissed biomechanics and fluid motion as confounds to be removed, potentially discarding valuable information about brain physiology. While this approach is intended to focus the analysis on signals of neuronal origin, it leaves unresolved the question of how ultra-slow fluctuations arise and organize across space at the macroscale.

In this study, we analyze resting-state fMRI recordings from 2177 scans from the Alzheimer’s Disease Neuroimaging Initiative (ADNI) repository, spanning the full spectrum of cognitive decline from cognitively normal (CN) aging through MCI to dementia (DEM)^41^. Avoiding any assumption regarding the origin of the fMRI signal fluctuations, we minimally preprocessed the data without removing CSF components. We use the term Brain-Ventricle (BraVe) rather than BOLD-CSF coupling to acknowledge that the only differentiating factor between signals is their anatomical location. Critically, this is not merely a terminological distinction: gray matter itself is substantially fluid-filled, with interstitial fluid occupying 15 to 20% of total brain volume and the brain consisting of approximately 80% water overall^42,43^. As a result, EPI signals in gray matter voxels are sensitive not only to blood oxygenation changes via neurovascular coupling, but also to interstitial fluid dynamics, perivascular flow, and tissue displacement, all of which can alter the EPI signal in the same way as CSF movement in ventricular voxels. The term BraVe therefore reflects a physically grounded agnosticism about signal origin: what differs between brain parenchyma and ventricular voxels is anatomy, without making assumptions regarding the origin of the signal. We extend the Leading Eigenvector Dynamics Analysis (LEiDA) algorithm^23,44^ beyond gray matter to identify recurrent modes of phase coupling across the entire brain volume including ventricular spaces, aiming to reveal hidden features of correlated signal fluctuations at the macroscale that are otherwise missed by segmented analysis approaches.

## 2. Results

### 2.1. BraVe mode occupancy tracks the progression of cognitive decline

Applying LEiDA to whole-brain fMRI signals, agnostic to signal origin, anatomical location, or participants’ clinical status, revealed 209 recurrent modes of phase alignment across a range of K-means clustering solutions (K = 2 to 20; see Methods and Supplementary Information for details of the LEiDA pipeline extension). Among these, modes whose occupancy differed significantly between diagnostic groups after conservative Bonferroni correction (α = 0.05/209 = 2.39 × 10^-4^) were retained and clustered by spatial similarity (cc > 0.65), yielding four representative BraVe modes that are the focus of the present report (**Figure 1**).

**Figure 1.**
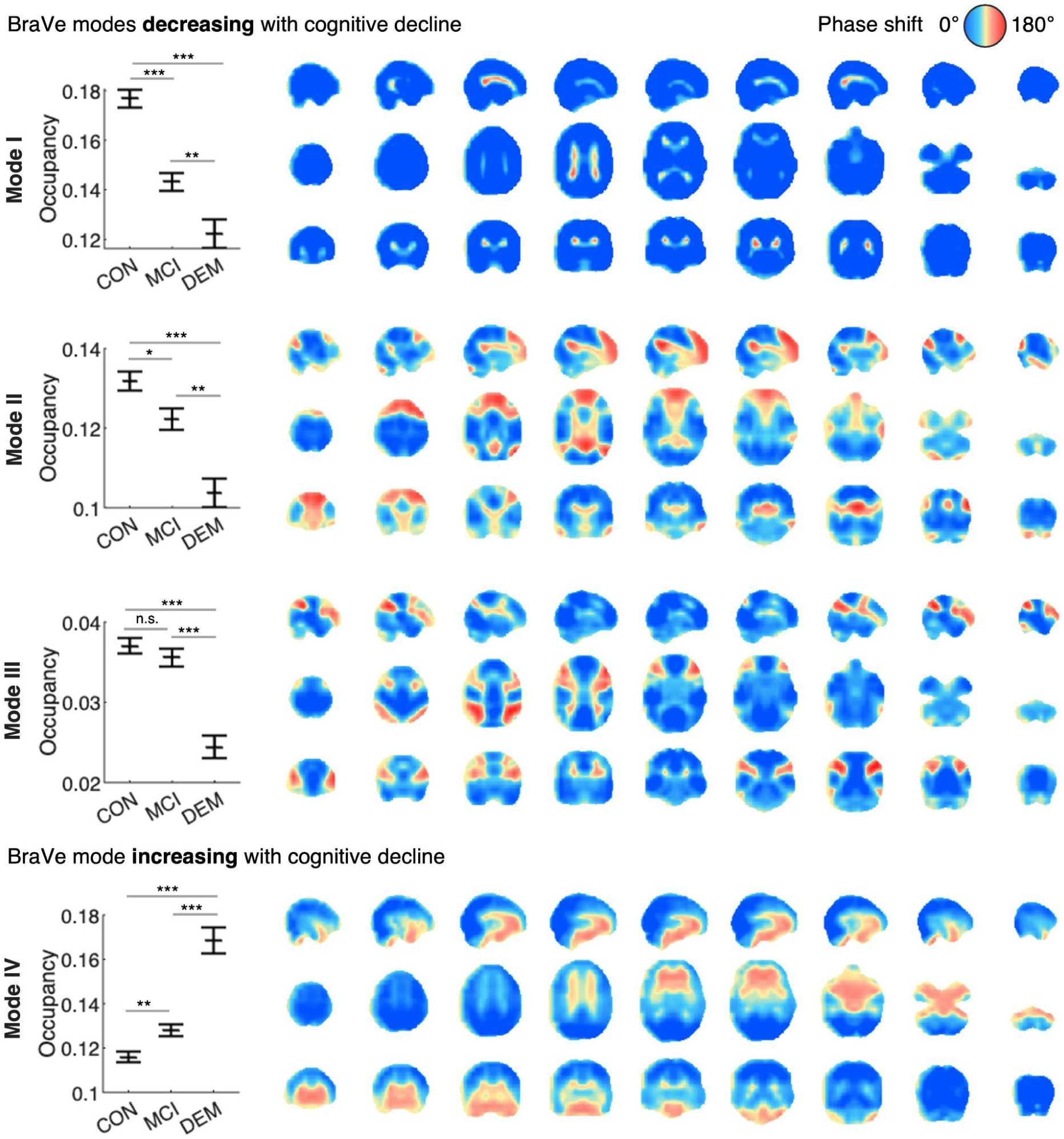
BraVe modes with occupancy increasing or decreasing with cognitive decline. (Left) Mean occupancy in each condition (Cognitively Normal (CN), Mild Cognitive Impairment (MCI) and Dementia (DEM)), with error bars representing the standard error of the mean (SEM) and asterisks indicating group differences from permutation testing (* p < 0.05, ** p < 0.001, *** p < 2.39 × 10^-4^). Modes I through III show progressively lower occupancy from CN to MCI and DEM, whereas Mode IV increases in occupancy. **(Right)** Each BraVe mode is represented by the average of all leading eigenvectors assigned to each mode. Each element in the eigenvector map corresponds to a brain voxel, in which blue voxels evolve within 90° (π/2) of the collective phase orientation, while red voxels exhibit a phase shift greater than 90°, indicating anti-phase behaviour relative to the collective orientation.

Occupancy, defined as the proportion of timepoints assigned to a given mode within each scan, was compared across 1045 CN, 831 MCI, and 301 DEM scans. CN individuals consistently showed the highest occupancy for Modes I, II, and III while DEM showed the lowest, with the reverse pattern for Mode IV.

Differences between CN and MCI were the most subtle, with significant effects surviving Bonferroni correction only for Mode I (p = 2.38 × 10^-8^, Hedge’s g = 0.29), while Mode II showed a nominal effect (p = 0.008, d = 0.12), Mode IV a weaker nominal effect (p = 0.001, g = 0.15), and Mode III showed no significant difference (p = 0.324, g = 0.05). Comparisons between MCI and DEM showed a complementary pattern, with Modes II, III, and IV reaching strict significance (Mode II: p = 1.9 × 10^-4^, g = 0.25; Mode III: p = 2.47 × 10^-8^, g = 0.38; Mode IV: p = 2.89 × 10^-8^, g = 0.46), while Mode I showed a nominal effect only (p = 0.004, g = 0.20). The CN versus DEM contrast yielded highly significant differences across all four modes, with effect sizes ranging from moderate to large (Mode I: p = 2.1 × 10^-8^, g = 0.46; Mode II: p = 2.43 × 10^-8^, g = 0.37; Mode III: p = 2.25 × 10^-8^, g = 0.42; Mode IV: p = 2.01 × 10^-8^, g = 0.63).

The spatial organization of each BraVe mode, visualized in sagittal, axial, and coronal slices **(Figure 1)**, reveals distinct patterns of phase alignment extending across the entire brain volume including ventricular spaces. Blue voxels oscillate within 90° degrees (or π/2 radians) of the collective phase orientation, indicating in-phase alignment with the dominant signal, while red voxels exhibit a phase shift greater than 90°, indicating anti-phase alignment. Crucially, in all four modes the ventricular voxels consistently appear in anti-phase with the majority of brain tissue voxels, justifying the term Brain-Ventricle (BraVe) coupling modes and confirming that the opposition between brain and ventricular signals is a recurring feature of resting-state fMRI dynamics varying across the cognitive spectrum.

To confirm that these findings are not an artefact of the longitudinal sampling structure, all group comparisons were replicated in two independent single-subject subsets (first scan per subject, N=880; last scan per subject, N=880). As shown in **Supplementary Table S3** and **Supplementary Figure S10**, the monotonic ordering of BraVe mode occupancy across diagnostic groups is preserved in both subsets, with effect sizes and significance levels consistent with the full sample. The CN vs DEM contrast remains highly significant across all four BraVe modes and both subsets (p < 0.00417 in all cases), confirming that the reported associations reflect genuine between-subject differences in brain dynamics across the cognitive decline spectrum.

### 2.2. BraVe modes reveal Resting-state Networks

Our results indicate the existence of functionally relevant modes of coupling extending beyond the dephasing of global brain signals from the ventricles. Given that the brain areas in which the signals temporarily dephase from the rest of the brain in modes II, III, and IV exhibit correlated activity at the cortical level, we turn to investigate how these BraVe modes relate to canonical RSNs. To do so, we calculated the spatial overlap between the set of dephasing voxels located in the cortex alone (**Figure 2b**) and the seven resting-state networks from a widely-used template^45^, defined on the same cortical voxels.

**Figure 2.**
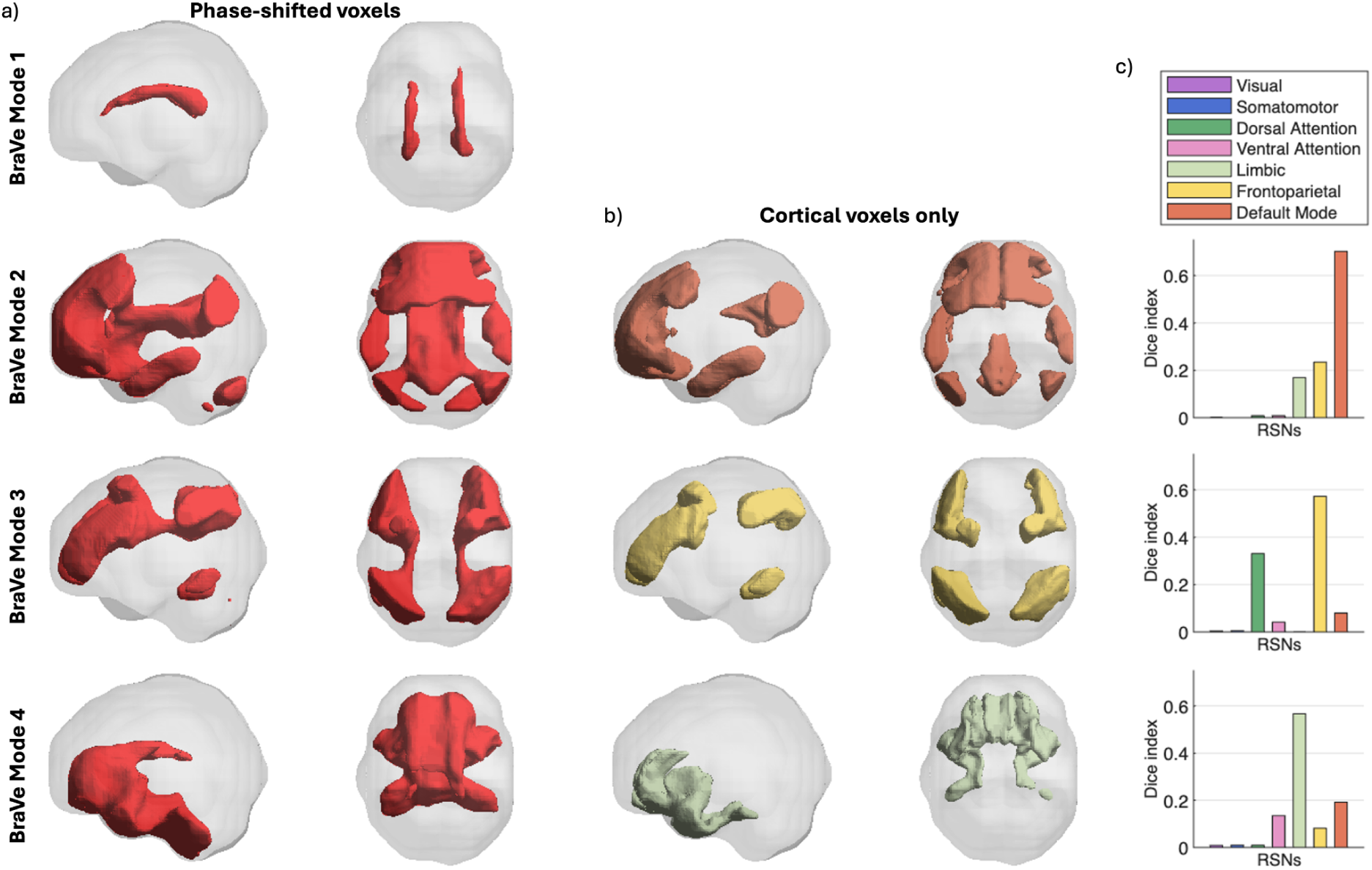
Cortical regions aligning with ventricle signals reveal canonical Resting State Networks (RSNs). **a)** 3D renderings of the 4 BraVe modes on a transparent brain mask, highlighting in red all voxels whose fMRI signals are phase-shifted by more than 90° relative to the rest of the brain. **b)** Same as a) but only including cortical voxels^43^. The colors correspond to the RSNs with maximal overlap. **c)** Dice coefficient between the binary mask obtained from b) and the masks of 7 canonical RSNs^43^.

Our analysis reveals that the BraVe modes shape functional networks at the cortical level, exhibiting a spatial overlap (Dice index>=0.55) with canonical RSNs (**Figure 2, bottom)**. Namely, BraVe mode II, decreasing with cognitive decline, overlaps strongly with the DMN (orange, diceindex =0.68). In particular, phase-shifts were larger in the superior frontal gyri, angular gyri, posterior cingulate gyri, and medial orbital frontal cortex, with additional voxels extending into the anterior cingulate gyri and middle and inferior temporal gyri, bilaterally (the complete list of gray matter areas defined with the AAL2 parcellation associated with each BraVe mode can be found in **Supplementary Figure S12).**

BraVe mode III, also occurring less frequently in cognitive impairment, shows strong overlap with the lateral Frontoparietal Network (FPN; yellow, Dice index = 0.58). Phase-shifts were largest in the inferior parietal lobules, inferior frontal gyri (triangular and opercular parts), middle frontal gyri, angular gyri, and superior parietal lobules bilaterally, with additional involvement of the supramarginal gyri, precentral gyri, and inferior temporal gyri. Reduced occupancy of BraVe modes II and III is consistent with previously reported reductions in static DMN and frontoparietal functional connectivity in cognitive decline, since higher mode occupancy directly increases the time-averaged correlation between the regions involved.

BraVe mode IV exhibits strong overlap with the limbic network (Dice index = 0.58). Phase-shifts were largest in the medial and orbital frontal cortex bilaterally (including the medial orbital frontal cortex, olfactory sulcus cortex, posterior and anterior orbital subdivisions, and rectus gyri), amygdala, temporal poles, anterior cingulate gyri, caudate nuclei, parahippocampal gyri, insula, putamen, and hippocampus bilaterally, with additional cerebellar involvement including the nodulus and lower vermis bilaterally.

### 2.3 Expression of BraVe modes over time and across subjects

Given that the clustering assigns each volume to one BraVe mode, it is possible to visualize how these modes are expressed at the instantaneous level in fMRI signals. In **Figure 3**, we show the brain signals from 3 participants matched for age and sex whose mode occupancy was representative of each cognitive condition (CN, MCI, and DEM). Background colors indicate which of the 4 BraVe modes was dominant at each time point, revealing that all modes recur intermittently throughout the scan. As illustrated in Figure 3, an occurrence of a given mode corresponds to periods when the fMRI signals in certain brain voxels either co-increase (red) or co-decrease (blue) with respect to the mean. Although instantaneous fMRI snapshots are inherently noisy, characteristic spatial patterns can be recognized: the anticorrelation between global brain signals and lateral ventricles during BraVe I, and the correlated increase or decrease of areas belonging to the DMN during BraVe II. Crucially, a mode can be expressed either positively or negatively (c.f. BraVe I at t=24 and t=155 for the same subject), a property that is captured by eigenvector-based methods but missed by amplitude-based approaches.

**Figure 3.**
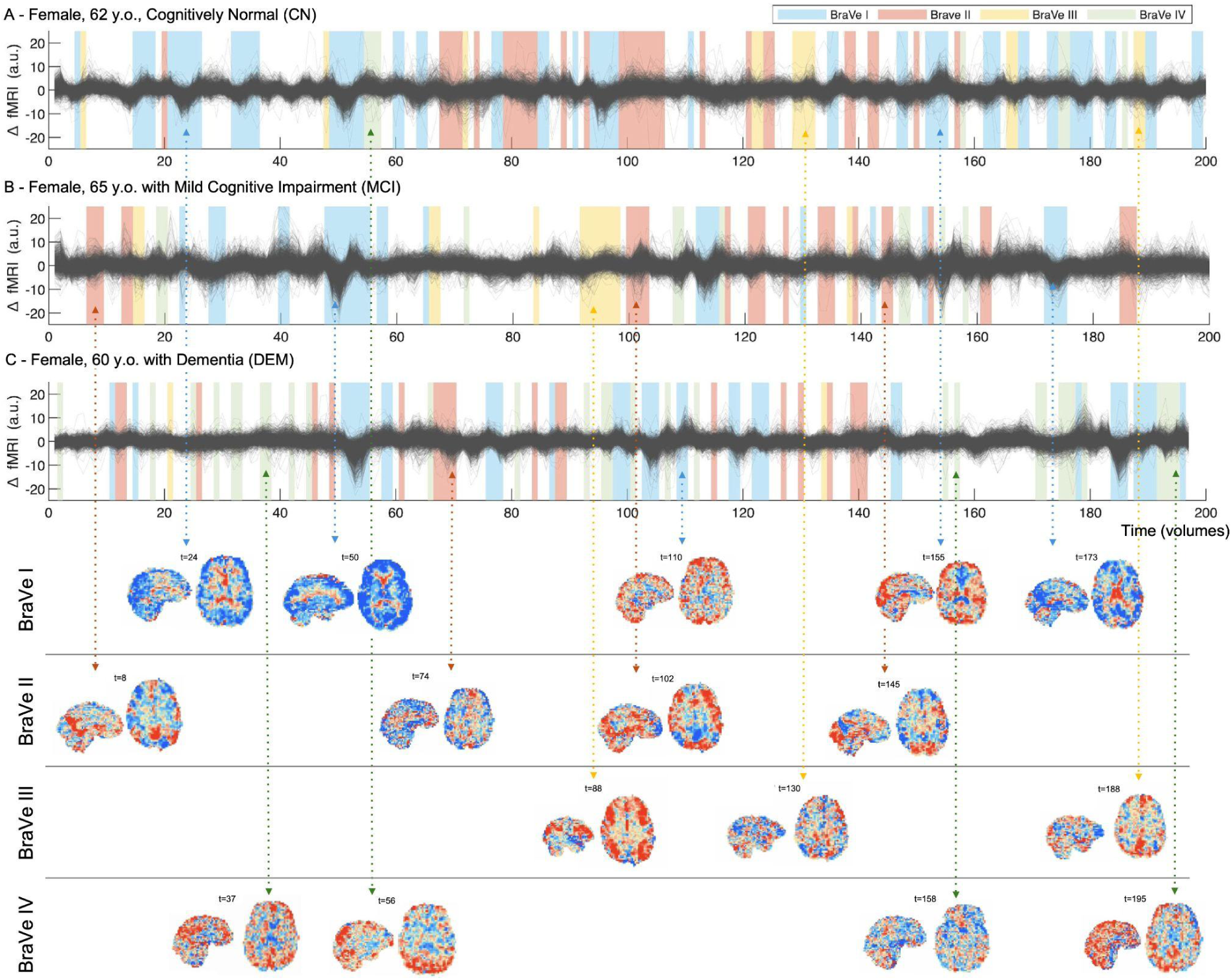
Brain dynamic states captured by BraVe modes in resting-state fMRI across cognitive decline stages. Panels A, B, and C show the preprocessed fMRI signal fluctuations (ΔfMRI in arbitrary units, a.u.) in all brain voxels over time for three representative female participants: a cognitively normal individual (CN, 62 years old, y.o.), a participant with Mild Cognitive Impairment (MCI, 65 y.o.), and a participant with Dementia (DEM, 60 y.o.). Background colors indicate the time points assigned to each of the 4 BraVe modes identified previously. Selected instantaneous fMRI volumes are displayed below for each mode, with dashed arrows linking each snapshot to its corresponding timepoint in the signal. Red and blue colors reflect positive and negative signal deviations from the voxelwise mean, respectively.

### 2.4. Occupancy of BraVe modes relates to cognition and disease biomarkers

We examined age-corrected partial correlations between BraVe mode occupancy and genetic risk, PET and CSF biomarkers of amyloid and tau pathology, and standardized cognitive and functional assessments across all scans (**Figure 4**, full details in **Supplementary Table S5**). The dominant pattern across all four modes was a robust and selective relationship with cognitive performance: between 16 and 27 variables per mode survived strict Bonferroni correction (p < 3.47 × 10^-4^), spanning global cognition, episodic memory, executive function, clinical severity, and everyday functioning. By contrast, molecular biomarkers were largely non-significant after age correction, with the notable exceptions of FDG-PET for Mode III and APOE ε4 allele count for Mode IV.

**Figure 4.**
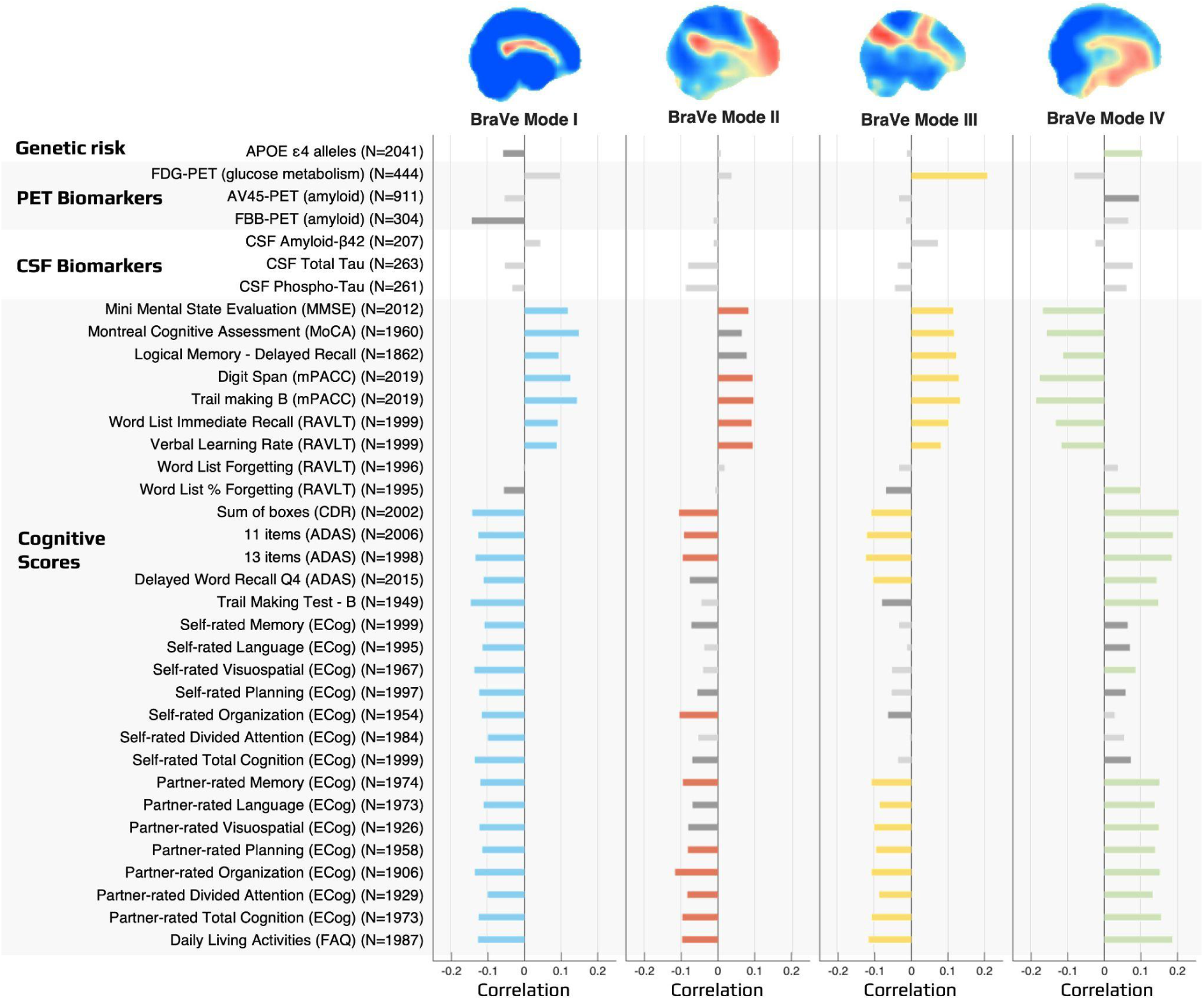
Age-corrected partial correlations between BraVe mode occupancy and genetic, PET biomarker, and psychometric variables. For each mode, bar plots represent the age-corrected partial Pearson correlation coefficient between mode occupancy and genetic risk factors, PET biomarkers, CSF biomarkers, and psychometric measures. Colored bars indicate associations surviving Bonferroni correction for 36 variables across 4 modes (p < 3.47 × 10^-4^). Dark gray bars indicate nominally significant associations (p < 0.005). Light gray bars indicate non-significant associations with p > 0.005. All correlations were computed as partial correlations with age included as a continuous covariate. Modified Preclinical Alzheimer Cognitive Composite (mPACC), Rey Auditory Verbal Learning Test (RAVLT), Clinical Dementia Rating (CDR), Alzheimer’s Disease Assessment Scale (ADAS), Everyday Cognition patient (ECog), Functional Activities Questionnaire (FAQ).

Within this overall pattern, each mode carried a distinct cognitive signature. Mode I showed the broadest profile, correlating significantly with virtually all cognitive and functional domains including both self- and partner-reported everyday cognition, episodic memory, and global screeners. Mode II captured a complementary but narrower set of associations, weighted toward partner-reported complaints and clinical severity. Mode III was the most selective, defined primarily by its association with FDG-PET metabolism (r = 0.20, p = 3.65×10^-5^, the strongest biomarker relationship found across all modes), alongside executive and memory scores. Mode IV showed the most severe clinical profile, correlating most strongly with CDR Sum of Boxes (r = 0.20, p = 1.13×10^-18^) and ADAS scores, and was the only mode whose occupancy linked to genetic AD risk: APOE ε4 allele count survived Bonferroni correction (r = 0.11, p = 1.30×10^-6^), with this association strengthening after age correction, indicating a genuine age-independent relationship between limbic entrainment and genetic vulnerability to AD.

Although individual effect sizes were modest, peaking around r = ±0.2, the mode-specific topology of these associations demonstrates that distinct BraVe coupling modes capture separable, biologically grounded dimensions of cognitive function and neurodegeneration.

### 2.5 Effects of age, education, and sex on BraVe mode occupancy

To characterise the independent contributions of demographic factors to BraVe mode occupancy, we performed a two-way ANCOVA with diagnostic condition and sex as factors and age and years of education as continuous covariates on site-harmonised occupancy values (**Supplementary Table S4**).

Diagnostic condition emerged as the dominant source of variance across all four modes (Mode I: p = 2.87 × 10^-16^; Mode II: p = 4.88 × 10^-5^; Mode III: p = 2.67 × 10^-7^; Mode IV: p = 1.56 × 10^-19^), with effect sizes substantially exceeding those of all other factors, and group differences persisting after controlling for age, education, and sex simultaneously.

Age showed a significant independent effect across all four modes (Mode I: p = 2.84 × 10^-5^; Mode II: p = 1.71 × 10^-10^; Mode III: p = 5.02 × 10^-7^; Mode IV: p = 1.70 × 10^-4^), indicating that advancing age contributes to occupancy changes beyond what is captured by diagnostic group alone. This is consistent with the well-documented trajectory of brain dynamics across the lifespan and suggests that the BraVe modes are sensitive to gradual age-related changes in brain-ventricle coupling independently of clinical diagnosis.

Education showed a significant independent effect specifically for BraVe Mode I (p = 3.67 × 10^-4^), with higher educational attainment associating with greater global synchronisation mode occupancy. This is consistent with the cognitive reserve hypothesis, whereby formal education builds resilience in large-scale brain dynamics that buffers against the effects of aging and neurodegeneration. Notably, education did not reach significance for Modes II, III, or IV (p = 0.515, p = 0.199, p = 0.118 respectively), suggesting that its protective influence is specifically expressed through the global synchronisation mode rather than through network-specific coupling patterns. This dissociation supports the interpretation that BraVe Mode I captures a functional signature of cognitive capacity that is sensitive to both reserve-building life experience and disease-related decline, while the network-specific modes reflect more targeted pathophysiological processes.

Sex showed a significant independent effect for BraVe Modes I and IV. Mode I showed higher occupancy in males (p = 5.33 × 10^-9^) without a significant difference in decline trajectory between sexes (Supplementary Figure S11), suggesting a baseline difference in global synchronisation that does not interact with disease progression. Mode IV showed both a significant sex main effect (p = 2.05 × 10^-6^) and a significant Condition × Sex interaction (p = 2.08 × 10^-3^), with females exhibiting markedly higher Mode IV occupancy as cognitive decline progresses, consistent with the stronger APOE ε4 penetrance reported in women^46^. Sex did not reach significance for Modes II and III (p = 0.037 and p = 0.178 respectively) after Bonferroni correction. Taken together, these results confirm that the group differences in BraVe mode occupancy cannot be attributed to demographic confounds, and that age, education, and sex each contribute meaningfully but independently and with substantially smaller effect sizes than diagnostic condition.

## 3. Discussion

This study reveals a set of recurring BraVe coupling modes in resting-state fMRI whose temporal occupancy tracks cognitive health across the full spectrum from normal aging to dementia. These findings bridge two fields that have largely developed in isolation: resting-state network research, focused on correlated slow fluctuations in gray matter, and brain-CSF coupling research^6,47^, which has concentrated almost exclusively on the global BOLD signal as a scalar marker of glymphatic function. Beyond confirming the global mode as a relevant marker of cognitive health, we show that other spatial modes overlapping with canonical resting-state networks also exhibit systematic relationships with cognitive status, disease biomarkers, and genetic risk, suggesting that the organisation of spontaneous brain activity into recurring spatial modes reflects the coupled physiology of neural tissue and ventricular fluid rather than neuronal communication alone. These results argue for an integrated framework in which resting-state dynamics and brain fluid regulation are understood as coupled aspects of a single system, with direct implications for fMRI-based biomarkers of neurodegeneration and brain clearance dysfunction. The precise physical mechanisms organising these modes remain to be elucidated.

### 3.1 Physical nature of resting-state dynamics and functional networks

The BraVe modes identified here challenge the conventional interpretation of resting-state fluctuations as driven solely by neuronal activity via neurovascular coupling. The EPI signal is equally sensitive to CSF flow and inflow effects and to spatial displacements at tissue-fluid interfaces, independently of neuronal activity. Indeed, lateral ventricle volume has been shown to oscillate in anti-phase with the global BOLD signal in standard fMRI acquisitions in the low-frequency range^48,49^.This sensitivity is not restricted to ventricular voxels: gray matter is substantially fluid-filled, with interstitial fluid occupying approximately 15 to 20% of total brain volume^42^ and the brain as a whole consisting of approximately 80% water^43^; EPI sequences at 3T have been shown to be directly sensitive to interstitial fluid volume in gray matter, independent of the BOLD signal^50^. The distinction between brain and ventricular signals in fMRI is therefore anatomical rather than mechanistic, and the conventional neurovascular framework neglects the fluid component present in every gray matter voxel, whose dynamics may contribute independently to the spatially organised fluctuations we observe.

The spatial organisation of RSNs presents further features difficult to explain under a purely neuronal framework: their bilateral symmetry with network poles consistently located over gyri and brain fissures, and their alignment with the geometric eigenmodes of the brain’s physical structure, suggest that anatomical and mechanical constraints play a fundamental organising role^51–53^. Ultrafast fMRI has revealed very low frequency CSF pulsations propagating with distinct spatiotemporal patterns through the brain, leading to the proposal that RSNs may reflect the spatial organisation of fluid-driven waves rather than axonal connectivity per se^11^. Failure of these glymphatic pulsation mechanisms has been proposed to precede protein accumulation in AD^11^, directly linking the dynamics captured in our BraVe modes to the neurodegenerative processes observed in our cohort. What our results add is that the proportion of time spent in specific BraVe coupling modes is a sensitive and clinically meaningful marker of cognitive health, independent of assumptions about its neuronal or non-neuronal origin.

### 3.2 BraVe modes as dimensional markers of cognitive function

Beyond group-level differences among CN, MCI, and DEM, BraVe mode occupancy shows continuous, graded relationships with cognitive performance across the full sample. All four modes show significant correlations with between 16 and 27 cognitive measures after strict Bonferroni correction, spanning global screening tools, episodic memory, executive function, functional independence, and informant-rated everyday cognition. The strongest single associations are for Mode IV, whose occupancy correlates with CDR Sum of Boxes (r=0.20, p=1.13×10^-18^), ADAS-11 (r=0.200, p=2.78×10^-18^), and inversely with the Trail Making Test B composite (r=-0.186, p=3.92×10^-16^), reflecting broader executive and functional decline. Mode III shows the strongest relationship with FDG-PET glucose metabolism (r = 0.20, p = 3.65×10^-5^), the only imaging biomarker surviving Bonferroni correction.

By contrast, molecular biomarkers show weaker statistical evidence: apart from APOE ε4 status for Mode IV, amyloid and tau measures do not survive correction, and CSF biomarkers are non-significant across all modes. The effect sizes for biomarker associations are nonetheless comparable to those for cognitive scores (r ∼ 0.09–0.20), suggesting that statistical power rather than biological irrelevance explains the difference, and larger multimodal datasets will be required to fully characterise the relationship between BraVe coupling and AD molecular pathology. This pattern suggests that BraVe modes track cognitive function as a continuous physiological dimension rather than serving as markers of AD-specific neuropathology, consistent with their potential relevance across neurological and psychiatric conditions beyond dementia.

### 3.3 Global synchronization and limbic entrainment as opposing poles of cognitive health

BraVe Mode I corresponds to the canonical global mode of brain-wide phase synchronization, a dynamical backbone of the healthy brain thought to support flexible information routing^23,54,55^. Its occupancy correlates robustly with global cognition (MMSE, MoCA), episodic memory, and executive function, and inversely with clinical severity (ADAS, CDR), establishing preserved global synchronization as a sensitive marker of cognitive health. Associations with FDG-PET and FBB-PET are in the expected direction but do not survive Bonferroni correction, consistent with the view that Mode I tracks domain-general cognitive function rather than AD-specific molecular pathology. The specific association of education with BraVe Mode I occupancy, and its absence for the remaining three modes, raises the possibility that time spent in global brain-ventricle synchronisation may be a more direct neural correlate of cognitive reserve than years of education alone. This would extend recent arguments that cognitive reserve is grounded not only in neuronal network organisation but also in neuroglial homeostatic mechanisms regulating brain fluid dynamics^56^, a hypothesis that prospective longitudinal studies combining BraVe analysis with glymphatic markers could test directly.

BraVe Mode IV represents the opposing dynamic: a state of limbic entrainment in which temporal regions, including the superior, middle, and inferior temporal gyri and fusiform cortex, shift into a specific phase relationship with the ventricles. Its occupancy increases progressively with cognitive decline and correlates with widespread functional impairment across objective and subjective domains. Given that amyloid accumulation in transmodal hubs disrupts local excitatory-inhibitory balance and destabilizes intrinsic oscillatory activity^57^, we propose that Mode IV reflects the macroscale consequence of this process, as local limbic perturbations propagate as phase drift and constrain brain dynamics within a regionally locked regime, progressively displacing the global synchronization captured by Mode I.

These findings extend prior observations linking APOE ε4 to both functional connectivity disruption in cingulate and medial prefrontal regions^58^ and reduced global gBOLD-CSF coupling^9^. BraVe Mode IV spatially refines both: its occupancy correlates with APOE ε4 allele count (r = 0.11, p =1.30×10^-6^), suggesting that genetic vulnerability to AD concentrates disruption within regions most susceptible to early amyloid accumulation.

### 3.4 Default mode erosion and the limits of frontoparietal compensation

BraVe Mode II shows marked overlap with the DMN, a canonical site of early amyloid deposition and metabolic vulnerability in AD^59^. The progressive decline in BraVe Mode II fractional occupancy across the neurodegenerative spectrum (CN > MCI > DEM), therefore, tracks the structural and functional erosion of this network in parallel with worsening global cognition and episodic memory. Notably, Aβ accumulation within DMN regions begins years before formal diagnosis^59^, and global glymphatic function, indexed by the coupling between global brain activity and CSF flow, has been shown to correlate with Aβ and tau markers at early disease stages^9^. This raises the possibility that reduced occupancy of BraVe Mode II reflects not only functional network deterioration but also impaired glymphatic clearance within these regions, with the spatial differentiation of BraVe modes potentially identifying which networks are most vulnerable to clearance-related pathology.

BraVe Mode III aligns with the lateral frontoparietal network, which provides flexible control over distributed systems to support goal-directed behaviour^60^. The nonlinear trajectory of Mode III occupancy, preserved from CN to MCI and then sharply declining at the transition to dementia, suggests that frontoparietal control remains relatively resilient during the prodromal phase. This stability is consistent with a compensatory response: as BraVe Mode II deteriorates, preservation of BraVe Mode III may buffer executive function and functional independence. The abrupt loss of Mode III occupancy at the transition to dementia may mark exhaustion of this frontoparietal reserve, indicating failure to recruit executive control networks to offset damage to core default mode systems. The specificity of this trajectory to the MCI-to-dementia transition, and its association with FDG-PET metabolism (r = 0.20, p = 3.65×10^-5^), the only imaging biomarker surviving Bonferroni correction across all modes, further underscores the metabolic sensitivity of frontoparietal function to advancing pathology.

### 3.5 Broader implications

An important practical implication concerns standard fMRI preprocessing. Ventricular CSF signals are routinely removed as nuisance regressors not only from ventricular voxels but also from gray matter signals, on the assumption that shared variance between CSF and cortical resting-state fMRI signals reflects physiological noise rather than meaningful dynamics. This preprocessing choice discards precisely the dynamics identified here as clinically informative, and may explain why prior studies restricted to gray matter have reported weaker associations between resting-state network occupancy and cognitive status. By retaining CSF-related variance and treating the entire brain volume as a unified signal space, the BraVe framework achieves statistical sensitivities not previously reported in this literature, with pairwise group differences reaching p < 2.01 × 10^-8^ and effect sizes up to g = 0.63. The framework can be applied retrospectively to any rsfMRI dataset in which CSF signals were preserved, opening existing archives to reanalysis and potentially revealing fluid-mediated markers of brain health across neurological and psychiatric conditions beyond cognitive decline.

### 3.6 Limitations

The relatively low temporal resolution (TR = 3 seconds) of the fMRI scans limits conclusions about the oscillatory or resonant nature of the observed phase locking. While recent studies with faster temporal resolution point to oscillatory and pulsatile processes at ultra-slow frequencies^6,11,14,61,62^, validating whether BraVe modes reflect genuine resonance in the strict sense requires higher temporal resolution than is available here. The space-time structure framework^63^ and the proposal that resting-state patterns reflect resonant eigenmodes of the brain’s physical structure ^51, 53^ provide relevant theoretical contexts for future work.

A second limitation is that we did not account for brain atrophy, which introduces discrepancies in spatial normalisation when using a young-adult MNI template in an elderly cohort (see ^64^ for a review). The current framework focused on macroscopic modes of phase alignment rather than precise anatomical locations, and the large sample size and conservative statistics reinforce the validity of the results. Future work should characterise BraVe mode occupancy across the healthy lifespan to establish whether the relationship with cognitive function observed here holds broadly across the adult age range, or emerges specifically as cognition begins to decline.

### 3.7 Conclusions

These findings invite a reconceptualisation of what resting-state fMRI measures: not merely the statistical fingerprint of neuronal communication, but the coupled physiology of brain tissue and ventricular fluid, whose organisation into recurring spatial modes tracks the brain’s capacity to sustain cognition and clear metabolic waste. BraVe mode occupancy emerges from this work as a sensitive, non-invasive, and biologically grounded marker of cognitive health, detectable with standard clinical fMRI acquisitions and interpretable within a framework that bridges network neuroscience and brain fluid dynamics. Beyond the clinical associations reported here, the BraVe framework reveals that the phase shifts underlying resting-state network dynamics are not arbitrary: cortical subnetworks shifting out of phase from the global brain signal are aligning with the ventricles, suggesting that RSN dynamics and brain-ventricle fluid coupling may be two perspectives on the same underlying physical process, one whose mechanistic basis, whether driven by CSF dynamics, cortical activity, or bi-directional resonance at infra-slow timescales, remains an important open question. As glymphatic research and resting-state fMRI converge on shared questions about brain clearance, resonance, and neurodegeneration, the BraVe framework offers a principled and tractable path toward integrated biomarkers of brain health across the lifespan.

## 4. Methods

### 4.1. Study population

The study included 880 unique subjects aged 50 to 90 years, acquired across 54 sites from the North American multicenter Alzheimer’s Disease Neuroimaging Initiative (ADNI; www.loni.ucla.edu/ADNI). Our study included all subjects from the ADNI2, ADNIGO, and ADNI3 phases who had resting-state functional magnetic resonance imaging (rsfMRI) scans and matching demographic and diagnostic information in the ADNIMERGE table (downloaded on February 6 2026). Our sample includes 301 fMRI scans from 103 participants diagnosed with Dementia (DEM) at the time of scan, 831 scans from 298 participants diagnosed with MCI, and 1045 scans from 479 cognitively normal (CN) participants (Baseline demographic characteristics of the 880 unique subjects, assessed at the time of first available scan, are summarised in Table 1, and further details can be found in Supplementary Information), totalling 2177 scans.

**Table 1.** Baseline characteristics of 880 unique subjects from ADNI included in this study, assessed at the time of first available scan. P-values were obtained using two-sample t-tests for continuous variables (age and education) and Fisher’s exact test for categorical variables (sex). Diagnostic group was assigned based on the clinical assessment at the time of the first scan. Statistically significant differences (p < 0.05) are marked in bold. CN = cognitively normal; DEM = dementia; MCI = mild cognitive impairment; M/F = male/female; SD = standard deviation. OR = Odds ratio.

| Variable | CN<br>(N=479) | MCI<br>(N=298) | DEM<br>(N=103) | CN vs MCI | CN vs DEM | MCI vs DEM |
| --- | --- | --- | --- | --- | --- | --- |
| Sex (M/F) | 192/287 | 166/132 | 57/46 | **$p=2.4 \times 10^{-5}$**<br>**OR=1.88** | **$p=0.006$**<br>**OR=1.85** | $p=1.0$<br>OR=0.99 |
| Age at baseline,<br>mean (SD) | 70.6 (6.6) | 71.8 (7.4) | 74.1 (7.2) | **$p=0.020$**<br>**$t=-2.32$** | **$p=1.7 \times 10^{-6}$**<br>**$t=-4.84$** | **$p=5.90 \times 10^{-3}$**<br>**$t=-2.76$** |
| Education years,<br>mean (SD) | 16.7 (2.3) | 16.0 (2.7) | 15.5 (2.5) | **$p=3.2 \times 10^{-4}$**<br>**$t=3.61$** | **$p=1.9 \times 10^{-6}$**<br>**$t=4.82$** | $p=0.069$<br>$t=1.82$ |

Where multiple scans were available for a given individual, all sessions were retained in the primary analysis. A formal longitudinal analysis tracking within-subject trajectories over time was not pursued here, as this would require accounting for the considerable heterogeneity in baseline clinical state, number of sessions, intervals between visits, and rate of clinical progression across individuals, which varies substantially in the ADNI cohort. Instead, each scan was treated as an independent observation characterised by its diagnostic label and cognitive scores at the time of acquisition, maximising statistical power for detecting associations between BraVe mode occupancy and the full spectrum of cognitive and biomarker measures. Diagnostic labels were assigned based on the clinical assessment conducted at the time of each scan, such that a single individual may contribute observations to more than one diagnostic group if their clinical status changed across visits. To confirm that the resulting group differences are not an artefact of the repeated-measures structure, all analyses were replicated in independent single-subject subsets restricted to the first and last available scan per individual (N=880 each; see Supplementary Table S3 and Supplementary Figure S10).

### 4.2. Cognitive scores and biomarkers

We considered all the variables available for these subjects in the ADNI database (ADNIMERGE table), namely genetic risk, in vivo PET and CSF biomarkers, and standardized cognitive and functional assessments obtained at the time of scan. The following variables available were used:

**Genetics:**

- APOE4: The å4 allele of apolipoprotein E, the strongest common genetic risk factor for AD^65^, is associated with increased amyloid deposition, earlier symptom onset, and accelerated disease progression^66^.

**PET biomarkers:**

- FDG: Fluorodeoxyglucose (^18^F) radiotracer reflecting cerebral glucose metabolism^67^; reduced uptake indicates synaptic dysfunction and neurodegeneration^68^.
- AV45: Florbetapir (^18^F) radiotracer quantifying fibrillar amyloid-β deposition in the cortex^69^, one of the defining pathological features of AD.
- FBB: Florbetaben (^18^F) radiotracer quantifying amyloid-β burden in the cerebral cortex, a hallmark pathological feature of AD^70^.

**CSF biomarkers:**

- Aâeta: Cerebrospinal fluid amyloid-â is generally interpreted as reflecting increased brain amyloid deposition and/or impaired amyloid clearance^71^.
- TAU: Total tau in CSF, a widely used marker of neuronal injury and axonal degeneration^72^;
- pTAU: Phosphorylated tau in CSF, a more specific indicator of tau pathology and neurofibrillary tangle burden^73,74^.

**Cognitive screening:**

- MMSE: Mini-Mental State Examination, a brief global cognitive screening tool that primarily captures orientation and general cognitive abilities^75^;
- MOCA: Montreal Cognitive Assessment, a sensitive screening instrument for MCI and early cognitive decline, with emphasis on executive function, attention, and memory^76^;
- Logical Memory - Delayed Recall: A verbal episodic memory task assessing delayed retention of story-based information;
- mPACC - Digit Span: A composite measure involving digit span, used to probe attention and WM within preclinical cognitive profiling;
- mPACC - Trial Making Test-B: A preclinical Alzheimer’s cognitive composite incorporating Trail Making Test-B, indexing set-shifting and executive control;
- RAVLT (Rey Auditory Verbal Learning Test^77^) - Immediate Recall: a measure of verbal learning and short-term episodic memory;
- RAVLT - Learning: The learning slope across repeated RAVLT trials, reflecting acquisition of verbal information over time;
- RAVLT - Forgetting: A delayed memory index quantifying loss of previously learned verbal material after a retention interval;
- RAVLT - % Forgetting: A normalized forgetting measure expressing retained versus lost verbal information after delay;
- CDR - Sum of Boxes: The Clinical Dementia Rating sum-of-boxes score, a composite staging measure of functional and cognitive impairment severity^78^;
- ADAS 11: Eleven-item Alzheimer’s Disease Assessment Scale-Cognition^79^ score, a broad measure of cognitive performance in AD trials;
- ADAS 13: Extended ADAS version with additional items, often used to improve sensitivity to early cognitive change;
- ADAS Q4 - Delayed Word Recall: The delayed recall subitem of ADAS-Cog, indexing episodic memory impairment;
- Trial Making Test-B: A standardized executive-function task measuring processing speed, cognitive flexibility, and set shifting^80^;
- ECog (Everyday Cognition^81^) Patient-rated (PT) or Informant-rated (SP) - Memory: memory domain, capturing perceived decline in daily memory function;
- ECog - Language: PT or SP everyday language functioning, reflecting subjective decline in word finding and communication;
- ECog - Visuospatial: PT or SP everyday visuospatial abilities, including navigation and spatial judgment;
- ECog - Planning: PT or SP executive/planning function in real-world activities;
- ECog - Organization: PT or SP ability to organize tasks and manage daily activities efficiently;
- ECog - Div. attention: PT or SP divided-attention capacity, reflecting multitasking and attentional control in daily life;
- ECog - Total: Global PT or SP everyday cognition score summarizing functional cognitive decline;
- FAQ (Functional Activities Questionnaire^82^), an informant-based measure of instrumental activities of daily living and functional independence.

### 4.3. MRI acquisition and data download

Resting-state fMRI scans were downloaded from the ADNI IDA repository across four phases (ADNI1, ADNI2, ADNIGO and ADNI3), acquired at 54 North-American sites using 3T scanners from multiple vendors (Siemens, GE, Philips) under five approved acquisition sequences: Axial rsfMRI (Eyes Open), Resting State fMRI, Axial MB rsfMRI (Eyes Open), Axial rsfMRI (EYES OPEN), and Extended Resting State fMRI, yielding 2,677 functional scans from 1,095 subjects. Acquisition parameters varied across sites and protocols, with full specifications available at https://adni.loni.usc.edu. Participants were instructed to keep their eyes open and remain awake during scanning. T1-weighted structural MRI scans were acquired in the same session using MPRAGE and equivalent sequences, and used for registration and tissue segmentation during preprocessing (see Supplementary Section S1 for full details).

### 4.4. fMRI preprocessing

Resting-state fMRI data were preprocessed using the Configurable Pipeline for the Analysis of Connectomes (C-PAC, v1.8.7), including slice timing correction, motion realignment to the mean functional volume using AFNI’s 3dvolreg, brain extraction using the niworkflows-ANTs method with the OASIS template, registration to MNI152 standard space (2 mm isotropic) using ANTs with a Rigid–Affine–SyN transformation strategy, spatial smoothing (5 mm FWHM), nuisance regression of the Friston 24 motion parameters and mean white matter signal, and temporal bandpass filtering (0.01–0.1 Hz) applied after nuisance regression. CSF signal was not included as a nuisance regressor in order to preserve CSF-related dynamics for subsequent analysis. After excluding scans that failed preprocessing, lacked demographic covariates, or originated from sites with fewer than 2 scans, the final dataset comprised 2,177 scans from 880 unique subjects across 54 acquisition sites. Full details of quality control steps, exclusion criteria, and pipeline configuration are provided in Supplementary Section S1.

### 4.5. fMRI data analysis

To analyze fMRI signals irrespective of their anatomical origin, each voxel inside a whole-brain mask was treated as a region of interest, rather than averaging signals within predefined parcels. A group-level brain mask was constructed by retaining voxels present in at least 95% of individual brain masks in 2mm isotropic MNI space. This mask was then resampled to an isotropic resolution of 10 mm using 3D linear interpolation, yielding N=1821 voxels covering the entire brain. The preprocessed fMRI volumes, already registered to MNI space, were downsampled from 2mm to 10mm using the same interpolation scheme, and the N=1821 voxels were treated as brain parcels for all subsequent analyses. This downsampling is justified given that signals were spatially smoothed with a 5mm FWHM Gaussian kernel during preprocessing, and larger voxels reduce between-subject variability in signal alignment.

To identify recurrent modes of whole-brain phase coupling and assess whether their occurrence varies with cognitive decline, we applied Leading Eigenvector Dynamics Analysis (LEiDA)^23^. For each voxel time series, the instantaneous phase θ(n,t) was extracted via the Hilbert transform. At each timepoint t (excluding the first and last volumes), the N×N matrix of instantaneous phase coherence was computed as cos(θ(n,t) − θ(p,t)) for all pairs of voxels n and p, and its leading eigenvector V1(t) was extracted. We note that, as a mathematical property of the cosine function, this phase coherence measure cannot distinguish between phase differences of 90° and 270°; eigenvector elements therefore span the interval [−1, 1], corresponding to mapped phase differences between 0° and 180°.

This 1×N vector captures the dominant mode of phase alignment across all N=1,821 brain voxels at that instant, with positive and negative components identifying voxels oscillating in phase and in anti-phase with the collective signal orientation, respectively. Applying this procedure across all 2,177 scans, each contributing one eigenvector per retained time point, yielded a total of 664,715 instantaneous eigenvectors, each providing a compact representation of the whole-brain phase-coupling pattern at a single time point. The average number of retained time points per scan was approximately 305, consistent with the typical rsfMRI acquisition length used across ADNI phases.

The 664,715 eigenvectors were then clustered into K discrete modes using k-means clustering with cosine distance, grouping timepoints that share a similar whole-brain phase-coupling pattern. Since the number of functionally relevant modes is not known a priori, K was varied from 2 to 20, and results were examined across the full partition range to ensure findings were not specific to a single choice of K. Stability was ensured by running 20 replicates per K and retaining the solution with the lowest total cluster-to-centroid distance. Each of the resulting 209 cluster centroids represents a recurrent Brain-Ventricle coupling mode, herein referred to as a BraVe mode^23^.

The clustering assigns a single cluster/mode to each time point by selecting the closest centroid. Using these cluster time courses, we calculated the Occupancy of each mode in each scan, defined as the number of time points assigned to a given state divided by the total number of time points (TRs) in that scan. The occupancies of each mode obtained with k ranging from 2 to 20 were calculated for each scan^23^.

### 4.6. Statistical analyses

Multi-site variability in fractional occupancy was removed using ComBat harmonization, preserving diagnostic group, age, education and sex as biological covariates of interest.

Welch’s t-test was preferred over Student’s t-test as it does not assume equality of variances across groups, making it appropriate for samples that differ in both size and variance. To avoid reliance on parametric assumptions about the sampling distribution of the test statistic, statistical significance was evaluated using a permutation procedure: group labels were randomly permuted 100,000 times and the Welch’s t-statistic was recomputed at each permutation to construct an empirical null distribution independently for each pairwise comparison. Two-sided p-values were derived as the proportion of permuted statistics whose absolute value equaled or exceeded the observed statistic. Effect sizes were quantified using Hedges’ g, which applies a sample correction to the pooled standard deviation estimate and is therefore preferred over Cohen’s d when group sizes are unequal, as is the case here.

To control the family-wise error rate across the large number of comparisons arising from testing multiple modes across multiple partition solutions, three progressively conservative significance thresholds were applied: α₁ = 0.05 (uncorrected), α₂ = 0.05/K (Bonferroni correction for the number of modes within each partition solution), and α₃ = 0.05/∑K = 0.05/209 = 2.39×10^-4^, which conservatively corrects for all modes tested across all partition solutions from K = 2 to K = 20 (∑K = 2 + 3 + … + 20 = 209), regardless of dependence between partitions. Only modes whose occupancy differences survived α₃ in at least one pairwise comparison were retained for further analysis.

Relationships between BraVe mode occupancy and cognitive scores, PET biomarkers, CSF biomarkers, and genetic risk factors were assessed using partial Pearson correlations across all scans, with age included as a continuous covariate to remove age-related variance independent of cognitive status. Correlations surviving Bonferroni correction for the number of variables tested (p < 3.47 × 10^-4^, correcting for 36 variables across 4 modes) are reported as significant.

To assess the contribution of sex to BraVe mode occupancy independently of demographic confounds, a two-way ANCOVA was performed with diagnostic condition and sex as categorical factors and age and years of education as continuous covariates. This analysis was motivated by a significant difference in educational attainment between males and females in this cohort (16.6 vs 15.9 years, p = 1.2 × 10^-11^), which could otherwise confound the observed sex effects. Group differences in BraVe mode occupancy were tested pairwise (CN vs MCI, CN vs DEM, MCI vs DEM) for each sex separately, and the Condition × Sex interaction term was used to assess whether the trajectory of occupancy change across diagnostic groups differed significantly between males and females. Bonferroni correction was applied across the four BraVe modes (p < 0.0125).

### 4.7 Visualization of cluster centroids

Each cluster centroid is a 1×N vector whose elements correspond to the N=1821 brain voxels at 10mm isotropic resolution. The sign of each element reflects the direction of each voxel’s phase relative to the collective phase orientation: voxels with positive values oscillate in phase with the dominant orientation (blue in figures). In contrast, voxels with negative values oscillate in anti-phase (red), as determined by the sign of the cosine of the phase difference. The smaller community, shown in red, identifies the brain areas shifting out of phase relative to the rest of the brain. For visualization, centroids were upsampled from 10mm to 2mm isotropic resolution using 3D linear interpolation.

### 4.8 Overlap with RSNs

The spatial overlap between each BraVe mode and seven canonical resting-state networks (RSNs) was assessed using the Yeo et al. cortical parcellation^45^. Cluster centroids were upsampled from 10mm^3^ to 2mm^3^ isotropic resolution, and a binary mask was derived by setting voxels with negative eigenvector values to 1 and all others to 0, considering cortical voxels only. The Dice coefficient was then computed between this binary mask and each of the seven RSN masks.

## 5. Acknowledgements

Data collection and sharing for this project were funded by the Alzheimer’s Disease Neuroimaging Initiative (ADNI) (National Institutes of Health Grant U01 AG024904) and DoD ADNI (Department of Defense award number W81XWH-12-2-0012). ADNI is funded by the National Institute on Aging, the National Institute of Biomedical Imaging and Bioengineering, and through generous contributions from the following: AbbVie, Alzheimer’s Association; Alzheimer’s Drug Discovery Foundation; Araclon Biotech; BioClinica, Inc.; Biogen; Bristol-Myers Squibb Company; CereSpir, Inc.; Cogstate; Eisai Inc.; Elan Pharmaceuticals, Inc.; Eli Lilly and Company; EuroImmun; F. Hoffmann-La Roche Ltd and its affiliated company Genentech, Inc.; Fujirebio; GE Healthcare; IXICO Ltd.; Janssen Alzheimer Immunotherapy Research & Development, LLC.; Johnson & Johnson Pharmaceutical Research & Development LLC.; Lumosity; Lundbeck; Merck & Co., Inc.; Meso Scale Diagnostics, LLC.; NeuroRx Research; Neurotrack Technologies; Novartis Pharmaceuticals Corporation; Pfizer Inc.; Piramal Imaging; Servier; Takeda Pharmaceutical Company; and Transition Therapeutics. The Canadian Institutes of Health Research is providing funds to support ADNI clinical sites in Canada. Private sector contributions are facilitated by the Foundation for the National Institutes of Health (www.fnih.org). The grantee organization is the Northern California Institute for Research and Education, and the study is coordinated by the Alzheimer’s Therapeutic Research Institute at the University of Southern California. ADNI data are disseminated by the Laboratory for Neuro Imaging at the University of Southern California.

The Center for Music in the Brain (MIB) is funded by the Danish National Research Foundation (project number DNRF117).

This work was initiated during FFC’s PhD and, during that period, was fully funded by POR Puglia FESR FSE 2014–2020 (CUP: H99121006630008).

The authors acknowledge funding from the Italian Ministry of Research (MUR) in the framework of the National Recovery and Resilience Plan (NRRP) under the complementary actions to the NRRP “Fit4MedRob” Grant. (PNC0000007, n. B53C22006960001) funded by “NextGenerationEU”.

This publication was also produced with the co-funding European Union - Next Generation EU, in the context of The National Recovery and Resilience Plan, Investment Partenariato Esteso PE8 “Conseguenze e sfide dell’invecchiamento”, Project Age-It (Ageing Well in an Ageing Society), CUP: B83C22004800006.

JC is supported by LARSyS through funding by the Portuguese Foundation for Science and Technology (FCT) (doi: 10.54499/LA/P/0083/2020, 10.54499/UIDP/50009/2020 and 10.54499/UIDB/50009/2020)

VM was supported by the FCT doctoral grant 2025.02255.BDANA.

## 6. CRediT Author Statement

**FFC:** Investigation, Formal analyses, Data curation, Writing - Original Draft, Writing - Review & Editing; **VM:** Formal analyses, Data curation, Methodology, Software, Writing - Review & Editing; **EB:** Conceptualization, Supervision, Writing - Review & Editing; **SN:** Writing - Review & Editing; **BT:** Writing - Review & Editing; **GL:** Supervision, Writing - Review & Editing; **JC:** Conceptualization, Investigation, Supervision, Methodology, Software, Formal analyses, Data Curation, Visualization, Writing - Original Draft, Writing - Review & Editing.

## 7. Data and code availability statement

All scripts and data used in this study will be published on a public repository upon acceptance of the manuscript. The 4 BraVe modes will be uploaded as NIFTI files in MNI space to NeuroVault.

## Bibliography

1. Stern, Y. Cognitive reserve in ageing and Alzheimer’s disease. Lancet Neurol. 11, 1006–1012 (2012).

2. Arrondo, P., Elía-Zudaire, Ó., Martí-Andrés, G., Fernández-Seara, M. A. & Riverol, M. Grey matter changes on brain MRI in subjective cognitive decline: a systematic review. Alzheimers Res. Ther. 14, 98 (2022).

3. Bennett, D. A. et al. Neuropathology of older persons without cognitive impairment from two community-based studies. Neurology 66, 1837–1844 (2006).

4. Iliff, J. J. et al. A paravascular pathway facilitates CSF flow through the brain parenchyma and the clearance of interstitial solutes, including amyloid β. Sci. Transl. Med. 4, 147ra111 (2012).

5. Rasmussen, M. K., Mestre, H. & Nedergaard, M. The glymphatic pathway in neurological disorders. Lancet Neurol. 17, 1016–1024 (2018).

6. Fultz, N. E. et al. Coupled electrophysiological, hemodynamic, and cerebrospinal fluid oscillations in human sleep. Science 366, 628–631 (2019).

7. Roefs, E. C. A., Eiling, I., de Bresser, J., van Osch, M. J. P. & Hirschler, L. BOLD-CSF dynamics assessed using real-time phase contrast CSF flow interleaved with cortical BOLD MRI. Fluids Barriers CNS 21, 107 (2024).

8. Lewis, L. D. The interconnected causes and consequences of sleep in the brain. Science 374, 564–568 (2021).

9. Han, F. et al. Reduced coupling between cerebrospinal fluid flow and global brain activity is linked to Alzheimer disease–related pathology. PLOS Biol. 19, e3001233 (2021).

10. Han, F. et al. Decoupling of Global Brain Activity and Cerebrospinal Fluid Flow in Parkinson’s Disease Cognitive Decline. Mov. Disord. Off. J. Mov. Disord. Soc. 36, 2066–2076 (2021).

11. Kiviniemi, V. et al. Ultra-fast magnetic resonance encephalography of physiological brain activity - Glymphatic pulsation mechanisms? J. Cereb. Blood Flow Metab. Off. J. Int. Soc. Cereb. Blood Flow Metab. 36, 1033–1045 (2016).

12. Gu, Y. et al. Brain Activity Fluctuations Propagate as Waves Traversing the Cortical Hierarchy. Cereb. Cortex 31, 3986–4005 (2021).

13. Chen, J. E. et al. Resting-state ‘physiological networks’. NeuroImage 213, 116707 (2020).

14. Cabral, J., Fernandes, F. F. & Shemesh, N. Intrinsic macroscale oscillatory modes driving long range functional connectivity in female rat brains detected by ultrafast fMRI. Nat. Commun. 14, 375 (2023).

15. Raut, R. V. et al. Global waves synchronize the brain’s functional systems with fluctuating arousal. Sci. Adv. 7, eabf2709 (2021).

16. Liu, X., Zhang, N., Chang, C. & Duyn, J. H. Co-activation patterns in resting-state fMRI signals. NeuroImage 180, 485–494 (2018).

17. Lv, H. et al. Resting-State Functional MRI: Everything That Nonexperts Have Always Wanted to Know. AJNR Am. J. Neuroradiol. 39, 1390–1399 (2018).

18. Fox, M. D. et al. The human brain is intrinsically organized into dynamic, anticorrelated functional networks. Proc. Natl. Acad. Sci. U. S. A. 102, 9673–9678 (2005).

19. Zhang, S.-P. et al. Frequency dependent whole-brain coactivation patterns analysis in Alzheimer’s disease. Front. Neurosci. 17, (2023).

20. Adhikari, M. H., Belloy, M. E., Van der Linden, A., Keliris, G. A. & Verhoye, M. Resting-State Co-activation Patterns as Promising Candidates for Prediction of Alzheimer’s Disease in Aged Mice. Front. Neural Circuits 14, (2021).

21. Young, P. N. E. et al. Imaging biomarkers in neurodegeneration: current and future practices. Alzheimers Res. Ther. 12, 49 (2020).

22. Teipel, S. et al. Multimodal imaging in Alzheimer’s disease: validity and usefulness for early detection. Lancet Neurol. 14, 1037–1053 (2015).

23. Cabral, J. et al. Cognitive performance in healthy older adults relates to spontaneous switching between states of functional connectivity during rest. Sci. Rep. 7, 5135 (2017).

24. Grieder, M., Wang, D. J. J., Dierks, T., Wahlund, L.-O. & Jann, K. Default Mode Network Complexity and Cognitive Decline in Mild Alzheimer’s Disease. Front. Neurosci. 12, 770 (2018).

25. Cha, J. et al. Functional alteration patterns of default mode networks: comparisons of normal aging, amnestic mild cognitive impairment and Alzheimer’s disease. Eur. J. Neurosci. 37, 1916–1924 (2013).

26. Ouchi, Y. & Kikuchi, M. A review of the default mode network in aging and dementia based on molecular imaging. Rev. Neurosci. 23, 263–268 (2012).

27. Ibrahim, B. et al. Diagnostic power of resting-state fMRI for detection of network connectivity in Alzheimer’s disease and mild cognitive impairment: A systematic review. Hum. Brain Mapp. 42, 2941–2968 (2021).

28. Brier, M. R. et al. Loss of intranetwork and internetwork resting state functional connections with Alzheimer’s disease progression. J. Neurosci. Off. J. Soc. Neurosci. 32, 8890–8899 (2012).

29. Gili, T. et al. Regional brain atrophy and functional disconnection across Alzheimer’s disease evolution. J. Neurol. Neurosurg. Psychiatry 82, 58–66 (2011).

30. Griffanti, L. et al. Effective artifact removal in resting state fMRI data improves detection of DMN functional connectivity alteration in Alzheimer’s disease. Front. Hum. Neurosci. 9, (2015).

31. Wang, Z. et al. Differentially disrupted functional connectivity of the subregions of the amygdala in Alzheimer’s disease. J. X-Ray Sci. Technol. 24, 329–342 (2016).

32. He, X. et al. Abnormal salience network in normal aging and in amnestic mild cognitive impairment and Alzheimer’s disease. Hum. Brain Mapp. 35, 3446–3464 (2014).

33. Agosta, F. et al. Resting state fMRI in Alzheimer’s disease: beyond the default mode network. Neurobiol. Aging 33, 1564–1578 (2012).

34. Zhao, Q., Lu, H., Metmer, H., Li, W. X. Y. & Lu, J. Evaluating functional connectivity of executive control network and frontoparietal network in Alzheimer’s disease. Brain Res. 1678, 262–272 (2018).

35. Zhao, Q., Sang, X., Metmer, H., Swati, Z. nawab N. K. & Lu, J. Functional segregation of executive control network and frontoparietal network in Alzheimer’s disease. Cortex 120, 36–48 (2019).

36. Sridharan, D., Levitin, D. J. & Menon, V. A critical role for the right fronto-insular cortex in switching between central-executive and default-mode networks. Proc. Natl. Acad. Sci. U. S. A. 105, 12569–12574 (2008).

37. Westlye, E. T., Lundervold, A., Rootwelt, H., Lundervold, A. J. & Westlye, L. T. Increased hippocampal default mode synchronization during rest in middle-aged and elderly APOE ε4 carriers: relationships with memory performance. J. Neurosci. Off. J. Soc. Neurosci. 31, 7775–7783 (2011).

38. Tang, F. et al. Differences Changes in Cerebellar Functional Connectivity Between Mild Cognitive Impairment and Alzheimer’s Disease: A Seed-Based Approach. Front. Neurol. 12, 645171 (2021).

39. Yao, H. et al. Decreased functional connectivity of the amygdala in Alzheimer’s disease revealed by resting-state fMRI. Eur. J. Radiol. 82, 1531–1538 (2013).

40. Zhou, B. et al. Impaired functional connectivity of the thalamus in Alzheimer’s disease and mild cognitive impairment: a resting-state fMRI study. Curr. Alzheimer Res. 10, 754–766 (2013).

41. Beckett, L. A. et al. The Alzheimer’s Disease Neuroimaging Initiative phase 2: Increasing the length, breadth, and depth of our understanding. Alzheimers Dement. J. Alzheimers Assoc. 11, 823–831 (2015).

42. Lei, Y., Han, H., Yuan, F., Javeed, A. & Zhao, Y. The brain interstitial system: Anatomy, modeling, in vivo measurement, and applications. Prog. Neurobiol. 157, 230–246 (2017).

43. MacAulay, N. Molecular mechanisms of brain water transport. Nat. Rev. Neurosci. 22, 326–344 (2021).

44. Cabral, J., Kringelbach, M. L. & Deco, G. Functional connectivity dynamically evolves on multiple time-scales over a static structural connectome: Models and mechanisms. NeuroImage 160, 84–96 (2017).

45. Yeo, B. T. T. et al. The organization of the human cerebral cortex estimated by intrinsic functional connectivity. J. Neurophysiol. 106, 1125–1165 (2011).

46. Altmann, A., Tian, L., Henderson, V. W., Greicius, M. D., & Alzheimer’s Disease Neuroimaging Initiative Investigators. Sex modifies the APOE-related risk of developing Alzheimer disease. Ann. Neurol. 75, 563–573 (2014).

47. Liu, X. Decoupling between brain activity and cerebrospinal fluid movement in neurological disorders. J. Magn. Reson. Imaging JMRI 60, 1743–1752 (2024).

48. Cheng, H. & Kennedy, D. P. Temporal fluctuation in lateral ventricle volume and its coupling with CSF inflow and global BOLD signal. Sci. Rep. 15, 30537 (2025).

49. Yang, H.-C. (Shawn), et al. Coupling between cerebrovascular oscillations and CSF flow fluctuations during wakefulness: An fMRI study. J. Cereb. Blood Flow Metab. 42, 1091–1103 (2022).

50. Bender, B. & Klose, U. Cerebrospinal fluid and interstitial fluid volume measurements in the human brain at 3T with EPI. Magn. Reson. Med. 61, 834–841 (2009).

51. Cabral, J. & Sun, H. Beyond networks: Functional connectivity as eigenmode resonance. Aperture Neuro 6, (2026).

52. Gonçalves, I., Oliveira, D., Rocha, C., Cabral, J. & Parente, M. Brain modes of resonance estimated by a biophysical multi-compartment finite elements model. Sci. Rep. 15, 33845 (2025).

53. Pang, J. C. et al. Geometric constraints on human brain function. Nature 618, 566–574 (2023).

54. Bolt, T. et al. Autonomic physiological coupling of the global fMRI signal. Nat. Neurosci. 28, 1327–1335 (2025).

55. Raut, R. V. et al. Arousal as a universal embedding for spatiotemporal brain dynamics. Nature 647, 454–461 (2025).

56. Verkhratsky, A. & Zorec, R. Neuroglia in cognitive reserve. Mol. Psychiatry 29, 3962–3967 (2024).

57. Giorgio, J., Adams, J. N., Maass, A., Jagust, W. & Breakspear, M. Amyloid induced hyperexcitability in default mode network drives medial temporal hyperactivity and early tau accumulation. Neuron 112, 676–686.e4 (2024).

58. McKenna, F., Koo, B.-B., Killiany, R., & Alzheimer’s Disease Neuroimaging Initiative. Comparison of ApoE-related brain connectivity differences in early MCI and normal aging populations: an fMRI study. Brain Imaging Behav. 10, 970–983 (2016).

59. Palmqvist, S. et al. Earliest accumulation of β-amyloid occurs within the default-mode network and concurrently affects brain connectivity. Nat. Commun. 8, 1214 (2017).

60. Marek, S. & Dosenbach, N. U. F. The frontoparietal network: function, electrophysiology, and importance of individual precision mapping. Dialogues Clin. Neurosci. 20, 133–140 (2018).

61. Dowdle, L. T. et al. Statistical power or more precise insights into neuro-temporal dynamics? Assessing the benefits of rapid temporal sampling in fMRI. Prog. Neurobiol. 207, 102171 (2021).

62. Palmer, W. C., Park, S. M. & Levendovszky, S. R. Brain state transition analysis using ultra-fast fMRI differentiates MCI from cognitively normal controls. Front. Neurosci. 16, 975305 (2022).

63. Petkoski, S. & Jirsa, V. K. Normalizing the brain connectome for communication through synchronization. Netw. Neurosci. 6, 722–744 (2022).

64. MacDonald, M. E. & Pike, G. B. MRI of healthy brain aging: A review. NMR Biomed. 34, e4564 (2021).

65. Liu, C.-C., Liu, C.-C., Kanekiyo, T., Xu, H. & Bu, G. Apolipoprotein E and Alzheimer disease: risk, mechanisms and therapy. Nat. Rev. Neurol. 9, 106–118 (2013).

66. Serrano-Pozo, A., Das, S. & Hyman, B. T. APOE and Alzheimer’s Disease: Advances in Genetics, Pathophysiology, and Therapeutic Approaches. Lancet Neurol. 20, 68–80 (2021).

67. Mosconi, L., Pupi, A. & De Leon, M. J. Brain glucose hypometabolism and oxidative stress in preclinical Alzheimer’s disease. Ann. N. Y. Acad. Sci. 1147, 180–195 (2008).

68. Bouter, C. et al. 18F-FDG-PET Detects Drastic Changes in Brain Metabolism in the Tg4–42 Model of Alzheimer’s Disease. Front. Aging Neurosci. 10, 425 (2019).

69. Palmqvist, S. et al. Detailed comparison of amyloid PET and CSF biomarkers for identifying early Alzheimer disease. Neurology 85, 1240–1249 (2015).

70. Sabri, O. et al. Florbetaben PET imaging to detect amyloid beta plaques in Alzheimer’s disease: phase 3 study. Alzheimers Dement. J. Alzheimers Assoc. 11, 964–974 (2015).

71. Li, Y. et al. Decreased CSF clearance and increased brain amyloid in Alzheimer’s disease. Fluids Barriers CNS 19, 21 (2022).

72. Schraen-Maschke, S. et al. Tau as a Biomarker of Neurodegenerative Diseases. Biomark. Med. 2, 363–384 (2008).

73. Blennow, K. & Hampel, H. CSF markers for incipient Alzheimer’s disease. Lancet Neurol. 2, 605–613 (2003).

74. Buerger, K. et al. CSF phosphorylated tau protein correlates with neocortical neurofibrillary pathology in Alzheimer’s disease. Brain J. Neurol. 129, 3035–3041 (2006).

75. Folstein, M. F., Folstein, S. E. & McHugh, P. R. ‘Mini-mental state’. A practical method for grading the cognitive state of patients for the clinician. J. Psychiatr. Res. 12, 189–198 (1975).

76. Nasreddine, Z. S., et al. The Montreal Cognitive Assessment, MoCA: a brief screening tool for mild cognitive impairment. J. Am. Geriatr. Soc. 53, 695–699 (2005).

77. Estévez-González, A., Kulisevsky, J., Boltes, A., Otermín, P. & García-Sánchez, C. Rey verbal learning test is a useful tool for differential diagnosis in the preclinical phase of Alzheimer’s disease: comparison with mild cognitive impairment and normal aging. Int. J. Geriatr. Psychiatry 18, 1021–1028 (2003).

78. Williams, M. M., Storandt, M., Roe, C. M. & Morris, J. C. Progression of Alzheimer’s disease as measured by Clinical Dementia Rating Sum of Boxes scores. Alzheimers Dement. 9, S39–S44 (2013).

79. Rosen, W. G., Mohs, R. C. & Davis, K. L. A new rating scale for Alzheimer’s disease. Am. J. Psychiatry 141, 1356–1364 (1984).

80. Reitan, R. M. The relation of the trail making test to organic brain damage. J. Consult. Psychol. 19, 393–394 (1955).

81. Farias, S. T. et al. The measurement of everyday cognition (ECog): Scale development and psychometric properties. Neuropsychology 22, 531–544 (2008).

82. Pfeffer, R. I., Kurosaki, T. T., Harrah, C. H., Jr., Chance, J. M. & Filos, S. Measurement of Functional Activities in Older Adults in the Community. J. Gerontol. 37, 323–329 (1982).

