## Supplementary Information for "Cognitive function linked to temporal occupancy of Brain-Ventricle (BraVe) modes"

#### Content:


- **S1- Methods and Data Curation**
- **S2 - Participant Demographics**
- **S3 - Supplementary Figures**
- **S4 - Table Scores vs Mode Occupancies**
- **References**

### S1. Detailed Methods description

#### S1.1 Data Curation and Preprocessing

##### 1.1.1 Dataset — Alzheimer's Disease Neuroimaging Initiative (ADNI)

Data used in the preparation of this article were obtained from the Alzheimer's Disease Neuroimaging Initiative (ADNI) database (adni.loni.usc.edu). The ADNI was launched in 2003 as a public-private partnership, led by Principal Investigator Michael W. Weiner, MD. The primary goal of ADNI has been to test whether serial magnetic resonance imaging (MRI), positron emission tomography (PET), other biological markers, and clinical and neuropsychological assessment can be combined to measure the progression of mild cognitive impairment (MCI) and early Alzheimer's disease (AD). Subjects were scanned across four phases of the initiative: ADNI1, ADNI2, ADNIGO, and ADNI3. At each visit, subjects were classified as Cognitively Normal (CN), Subjective Memory Complaints (SMC), Early MCI (EMCI), Late MCI (LMCI), or Alzheimer's Disease (AD/Dementia). For all analyses, EMCI, LMCI and MCI were treated as a single MCI group, AD and Dementia as a single DEM group, and SMC was grouped with CN, as SMC subjects are cognitively normal by objective neuropsychological measures.

##### 1.1.2 Image Acquisition and Data Download

Resting-state functional MRI (rs-fMRI) scans were downloaded from the ADNI IDA repository using the following resting-state acquisition sequences: *Axial rsfMRI (Eyes Open)*, *Resting State fMRI*, *Axial MB rsfMRI (Eyes Open)*, *Axial rsfMRI (EYES OPEN)*, and *Extended Resting State fMRI*, yielding **1,095 subjects and 2,677 functional scans** across all ADNI phases. T1-weighted structural MRI scans were also downloaded using MPRAGE and equivalent sequences, yielding **1,099 subjects and 2,687 scans**, used for registration and tissue segmentation during preprocessing.

##### 1.1.3 BIDS Conversion

All downloaded scans were converted to the Brain Imaging Data Structure (BIDS) format (Gorgolewski et al., 2016) using *dcm2niix* for DICOM-to-NIfTI conversion and custom scripts for metadata organization. During conversion, **305 scans were identified as corrupted** (unreadable DICOM files, incomplete acquisitions, or missing metadata) and were excluded, resulting in **2,372 BIDS-compliant functional scans**.

##### 1.1.4 fMRI Preprocessing

Functional MRI data were preprocessed using the **Configurable Pipeline for the Analysis of Connectomes (C-PAC, v1.8.7)** with a pipeline configuration that includes the following preprocessing steps: (1) slice timing correction; (2) motion realignment using AFNI 3dvolreg with the mean functional image as reference; (3) brain extraction using the niworkflows-ANTs method with the OASIS template; (4) registration to MNI152 standard space (2mm isotropic) using Advanced Normalization Tools (ANTs) with a sequential Rigid–Affine–SyN transformation strategy; (5) spatial smoothing with a 5mm FWHM Gaussian kernel; (6) nuisance signal regression including the Friston 24 motion parameters (6 rigid-body parameters, their temporal derivatives, and their squares), mean white matter signal, and second-degree polynomial detrending; (7) temporal bandpass filtering (0.01-0.1 Hz). Scans that failed preprocessing due to insufficient timepoints or processing errors were excluded (**n=16**), resulting in **2,356 preprocessed functional scans**. The 16 excluded scans were: 002S0413/ses-005, 002S0685/ses-002, 002S1261/ses-004, 012S4026/ses-003, 013S4268/ses-005, 027S6733/ses-002, 027S6788/ses-002, 027S6842/ses-002, 031S4721/ses-004, 053S4578/ses-006, 053S4661/ses-005, 127S6173/ses-002, 130S4417/ses-006, 130S4883/ses-001, 136S4517/ses-002, 941S6499/ses-002.

##### 1.1.5 Quality Control and ADNIMERGE Alignment

Clinical scores, demographic variables, and diagnostic labels were obtained from the ADNIMERGE dataset (version 06/02/2026). All 2,356 preprocessed scans were successfully matched to ADNIMERGE records by subject identifier (PTID) and session identifier. Scans were excluded if they had missing values in acquisition site, age at scan, or sex (**N=177**), as these covariates are required for ComBat harmonization. An additional **2 scans** from sites with fewer than 2 scans were excluded. The final quality-controlled dataset comprised **2,177 scans from 880 unique subjects**, which was used for all subsequent steps.

| Step | Scans removed | Total scans |
|---|---|---|
| Downloaded from ADNI | —- | 2,687 |
| BIDS conversion | 305 | 2,372 |
| C-PAC preprocessed output | 16 | 2,356 |
| ADNIMERGE alignment | 0 | 2,356 |
| Missing age, sex, or site | 177 | 2,179 |
| Sites with < 2 scans | 2 | 2,177 |

**Supplementary Table S1. Quality control steps and scan counts.**

##### 1.1.6 Brain Voxels of Interest

During functional preprocessing, C-PAC generated a brain mask for each individual scan using the *Anatomical_Refined* method, which dilates the subject's anatomical brain mask and applies it to the functional data; each mask was subsequently brought into MNI152 2 mm isotropic standard space via the same registration warp used for the functional images. A group-level brain mask was then defined by averaging the individual MNI-space masks from the first 606 preprocessed scans and retaining only voxels present in at least 95% of scans. This 95% group mask was subsequently resampled to an isotropic resolution of 10 mm3, yielding a reduced set of N=1821 voxels, which were treated as parcels for all subsequent analyses.

##### 1.1.7 Leading Eigenvector Dynamics Analysis (LEiDA)

Brain dynamics were characterized using Leading Eigenvector Dynamics Analysis (LEiDA; Cabral et al., 2017). For each time point $t$ of each scan, the instantaneous phase was computed via the Hilbert transform of the bandpass-filtered signal at each brain voxel N. The NxN instantaneous phase coherence matrix *dFC(t)* was computed as the cosine of the phase difference between all pairs of voxels, and its 1xN leading eigenvector *V1(t)* was extracted for all volumes (except for the first and last volumes in each scan), yielding a total of **664715 eigenvectors** across the 2,177 scans with a mean length of 305 volumes.

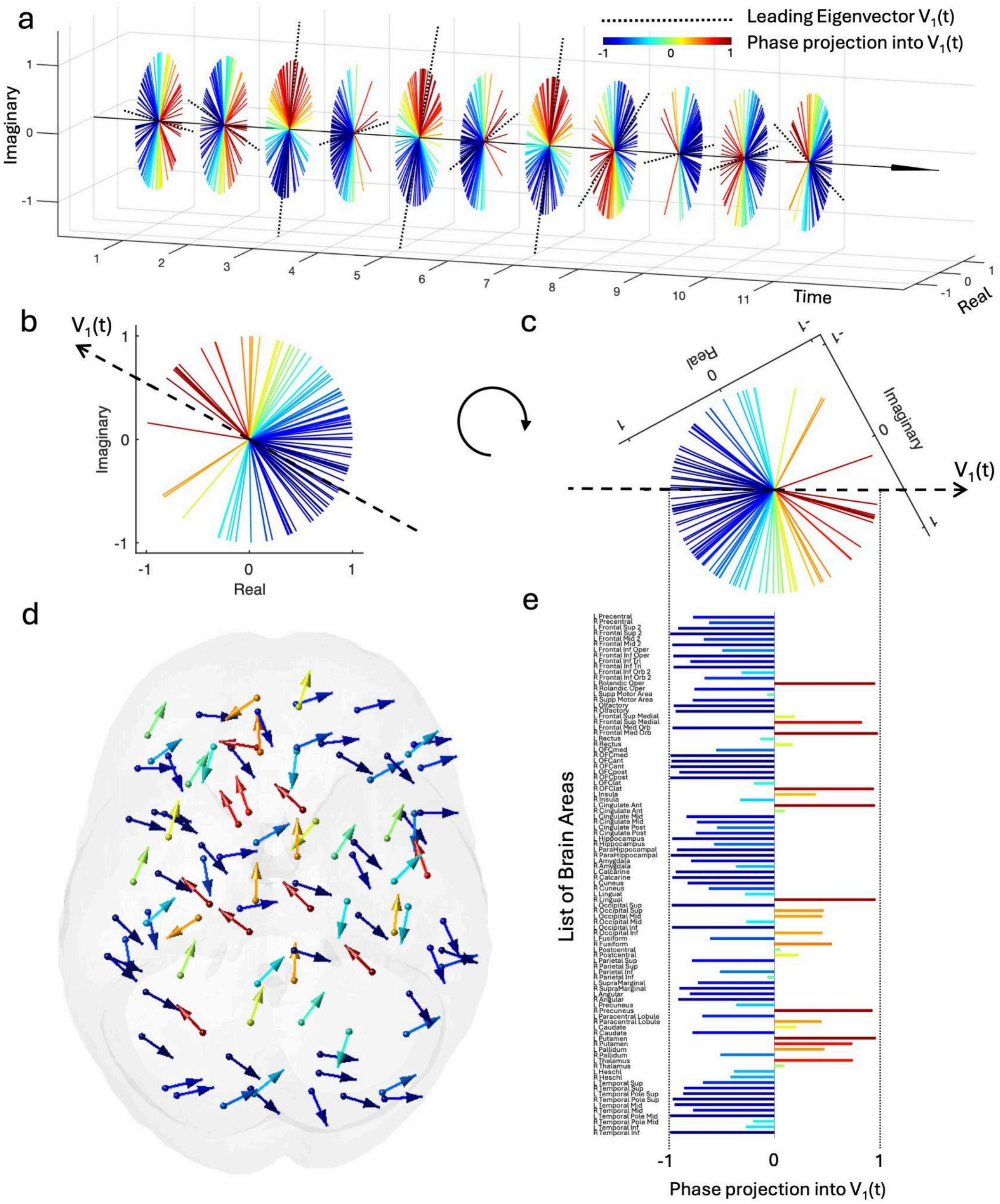


**Supplementary Figure S1. Illustration of the pattern captured by the leading eigenvector of phase coherence at each time point. (a)** After obtaining the signal phases in each parcel/voxel using the Hilbert transform, the leading eigenvector of the phase coherence matrix is obtained for each time point. The orientation of the leading eigenvector, $V_1(t)$, is represented as a black dashed line, and the phase of each parcel is colored according to its projection into $V_1(t)$. **(b)** At each time point, the signal phase in each parcel is represented in the complex plane. Given that the direction of the eigenvector is arbitrary ($V_1(t) = -V_1(t)$), we establish the convention that the majority of elements has negative sign (blue), such that only the parcels whose signals shift by more than $\pi/2$ with respect to the main orientation have positive values (red). **(c)** Considering the projection into the leading eigenvector instead of the Real axis corresponds to an axis rotation. **(d)** The instantaneous signal phases shown in (b) are represented as arrows originating from the center of each brain parcel. **(e)** Each element in the 1xN leading eigenvector corresponds to a brain area, and the values correspond to their projection into the axis defined by $V_1(t)$.

##### 1.1.8 K-means Clustering

K-means clustering was applied to all **664715** eigenvectors to identify recurrent whole-brain modes of phase alignment. Clustering was performed for K values from 2 to 20, with 20 replicates per K, ensuring stable results. Each brain mode is represented by a centroid vector, which corresponds to the mean of all the eigenvectors assigned to that cluster.

##### 1.1.9 Fractional Occupancy

For each scan and each K solution (k=2 to k=20), **fractional occupancy** P_k was computed as the proportion of timepoints assigned to each state k, yielding a vector of k values per scan. This measure quantifies the time the brain spends in each coupling mode and serves as the primary feature used in all subsequent statistical analyses.

##### 1.1.10 ComBat Harmonization of Fractional Occupancy

Prior to statistical analysis, multi-site and multi-protocol variability in fractional occupancy values were removed using **ComBat harmonization** (Johnson et al., 2007; Fortin et al., 2018), as implemented in the MATLAB ComBatHarmonization toolbox (github.com/Jfortin1/ComBatHarmonization). ComBat applies an empirical Bayes framework to estimate and remove additive and multiplicative batch effects, here defined as acquisition site, while preserving the biological variability of interest. The 2,177 scans originated from **54 acquisition sites** across the ADNI phases. ComBat was applied separately to the fractional occupancies at each K solution (k=1 to k=20). The biological covariates preserved during harmonization included: diagnostic group (CN/SMC=0, MCI=1, AD/Dementia=2), age at scan, sex and education. Parametric adjustment was used. SMC subjects were grouped with CN for both the ComBat covariate model and all statistical analyses, as SMC subjects are cognitively normal by objective neuropsychological criteria.

##### 1.1.11 Final Cohort

The final dataset comprised **2,177 scans from 880 unique subjects** across ADNI1, ADNI2, ADNIGO, and ADNI3, collected at 54 sites. Scans were acquired using three collection protocols: ADNI2 (n=763), ADNIGO (n=54), and ADNI3 (n=1,360), from subjects originally recruited across all four ADNI phases (ADNI1: n=144; ADNI2: n=1,062; ADNIGO: n=165; ADNI3: n=806). The diagnostic composition at time of scan is shown in Table 2.

| Diagnostic Group | N scans | % of total | Notes |
|---|---|---|---|
| CN + SMC | 1,045 | 48.0% | SMC grouped with CN |
| MCI (EMCI + LMCI + MCI) | 831 | 38.2% | |
| AD + Dementia | 301 | 13.8% | |
| Total | 2,177 | 100% | 880 unique subjects, 54 sites |

Supplementary Table S2. Final cohort composition after all preprocessing and quality control steps.

### S2 - Participant Demographics

ADNI inclusion and exclusion criteria are detailed at http://www.adni-info.org procedures manual.

Sex distribution at baseline differed significantly across diagnostic groups. Given the categorical nature of this variable, pairwise comparisons were conducted using Fisher's exact test. In the CN group, females were overrepresented ($n$ = 287, 59.92%) relative to males ($n$ = 192, 40.08%). This pattern was reversed in the clinical groups: in MCI, females accounted for 132 (44.30%) and males for 166 (55.70%), whereas in DEM, females accounted for 46 (44.66%) and males for 57 (55.34%). The CN versus MCI comparison revealed a highly significant difference ($p = 2.42\times10^{-5}$; odds ratio = 1.88), indicating that the odds of being female were nearly twofold higher in the control group than in the MCI group. A similar pattern was observed for CN versus DEM ($p = 5.90\times10^{-3}$, odds ratio = 1.85), again reflecting a higher proportion of females among CN. In contrast, no difference emerged between MCI and DEM ($p$ = 1.00; odds ratio = 0.99), indicating virtually identical sex distributions between the two groups.

Age at baseline showed a graded and statistically robust increase across diagnostic groups. In the comparison between CN and MCI, a significant difference was observed ($t$ = -2.32, $p$ = 0.020), indicating that MCI were, on average, older than CN even at baseline. This effect was markedly amplified when comparing CN and DEM, with a substantially larger effect size ($t$ = -4.84, $p = 1.70\times10^{-6}$), supporting a pronounced age gap between these groups. The comparison between MCI and DEM further confirmed this progression, with DEM being significantly older than MCI ($t$ = -2.76, $p$ = 0.006). Mean age increased from 70.57 ± 6.57 years in CN (n=479) to 71.76 ± 7.42 years in MCI (n=298), and 74.08 ± 7.18 years in DEM (n=103).

Lastly, years of education at baseline exhibited an inverse pattern, decreasing with increasing disease severity. The comparison between CN and MCI revealed a highly significant difference ($t$ = 3.61, $p = 3.20\times10^{-4}$), with the control group showing higher educational attainment. This difference became more pronounced in the CN vs. DEM comparison ($t$ = 4.82, $p = 1.86\times10^{-6}$), further indicating that CN had substantially higher levels of education than DEM. In contrast, the comparison between MCI and DEM did not reach conventional statistical significance ($t$ = 1.82, $p$ = 0.069), although a trend toward higher education in MCI was observed. Mean years of education were 16.68 ± 2.29 in CN, 16.02 ± 2.74 in MCI, and 15.46 ± 2.52 in DEM.

### S3 - Supplementary Figures

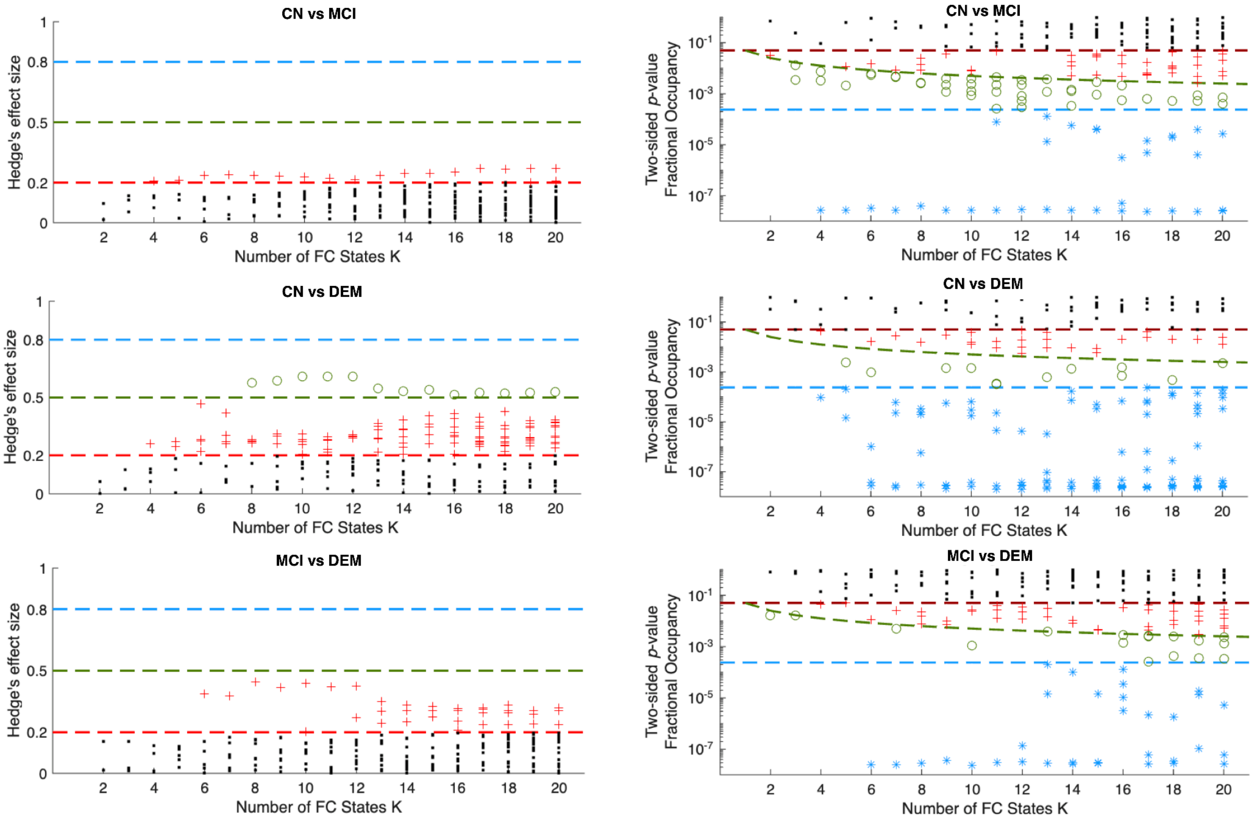


**Supplementary Figure S2**. **Statistical analysis of mode occupancies between cognitive status groups.** The modes obtained with K-means clustering with K varying between 2 and 20, were compared across the 3 stages of cognitive decline. (**Left**) Effect sizes (Hedge's g) represent the magnitude of differences across conditions. Red indicates a small effect size (0.2 < g < 0.49), green a medium effect size (0.5 < g < 0.79), and blue a large effect size (g > 0.8). (**Right**) p-values from the permutation test comparing occupancies between condition pairs. Red represents $p_{perm} < 0.05$, green represents $p_{perm} < 0.05/K$, and blue represents $p_{perm} < 2.39 \times 10\text{-}4$.

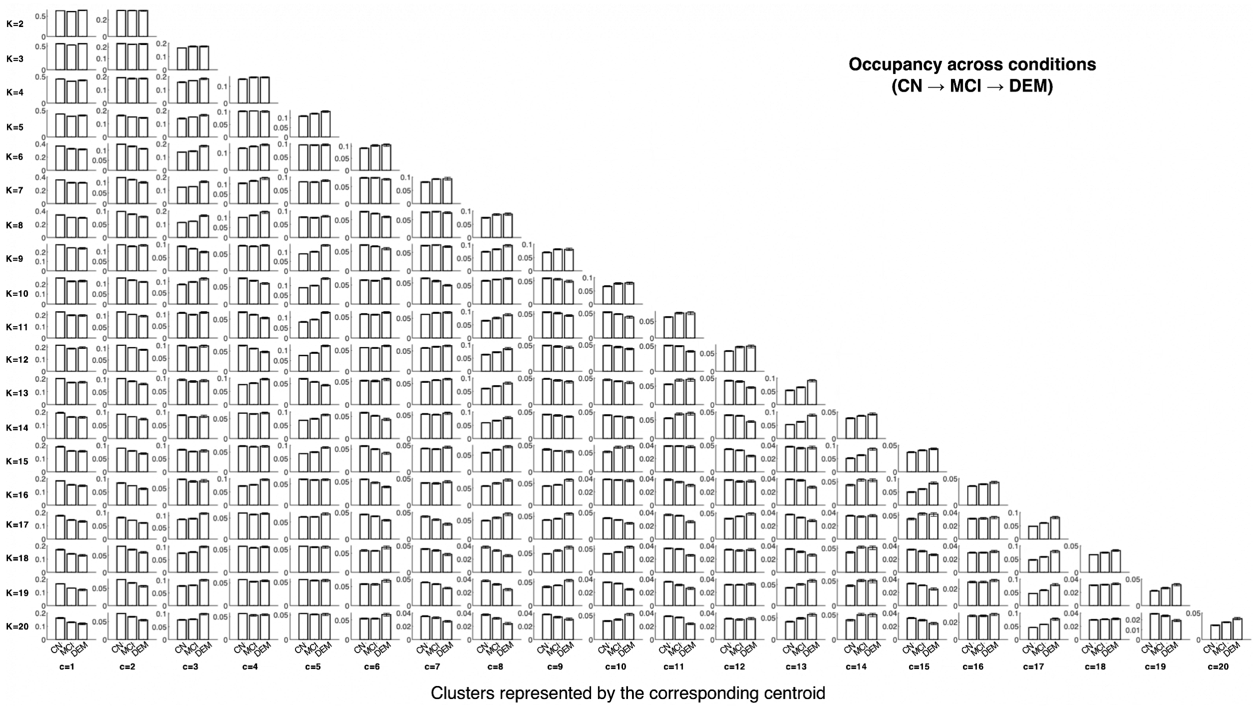


**Supplementary Figure S3 - Occupancy of Phase Coupling Modes in each cognitive status group.** For each K (2 to 20), bar plots show occupancy in each condition (from left to right: CN, MCI, DEM).

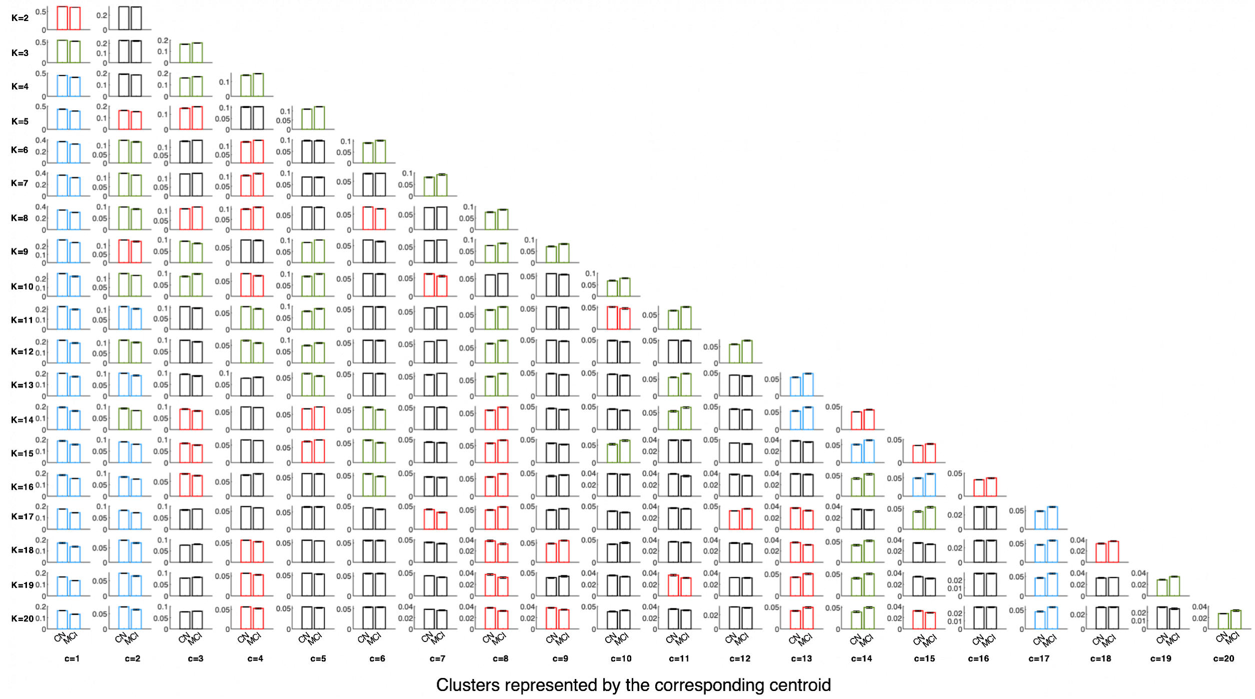


**Supplementary Figure S4 - Changes in Mode Occupancy between Cognitively Normal (CN) and Mild Cognitive Impairment (MCI) participants.** Bar plots showing significant differences in occupancy between CN and MCI. Red bar plots indicate $p_{perm} < 0.05$, below the significance threshold, albeit with a risk of false positives; green bar plots indicate $p_{perm} =< 0.05/K$, while blue bar plots indicate $p_{perm} < 2.39 \times 10^{-4}$.

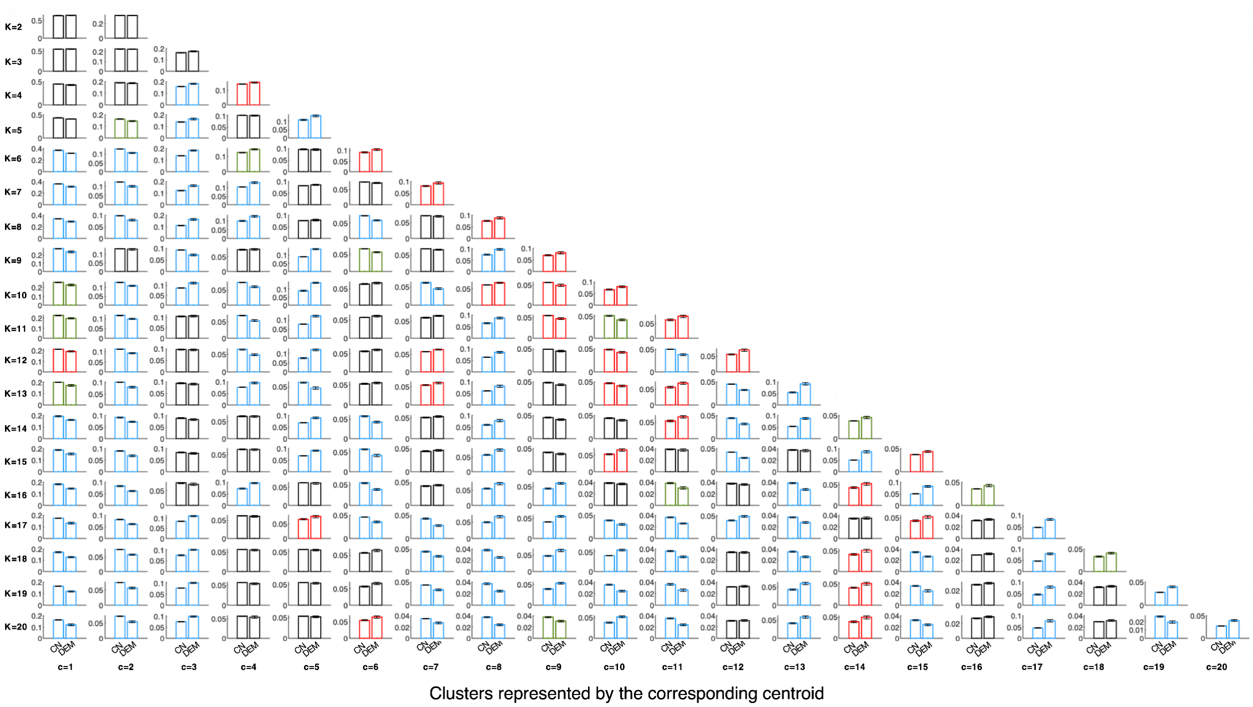


**Supplementary Figure S5 - Changes in Mode Occupancy between Cognitively Normal (CN) and participants with Dementia (DEM).** Barplots showing occupancy in CN and DEM, highlighting differences in red for $p_{perm} < 0.05$, green for $p_{perm} < 0.05/20$, and blue for $p_{perm} < 2.39 \times 10^{-4}$.

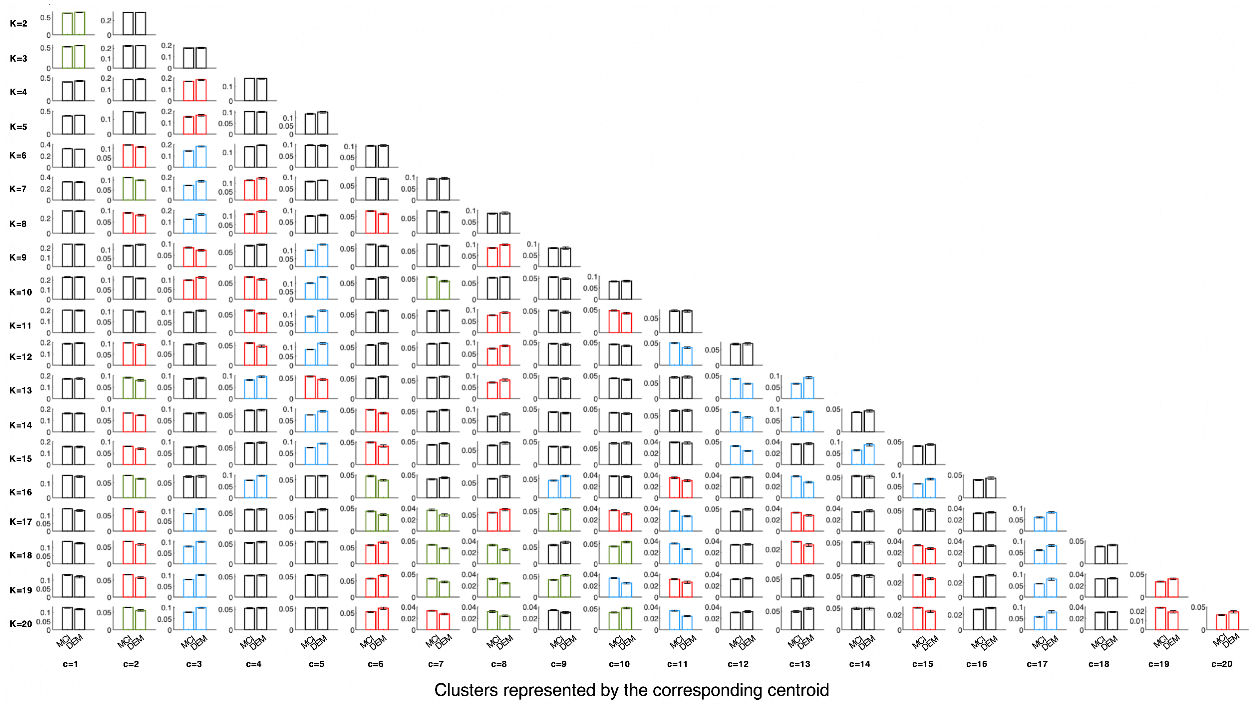


**Supplementary Figure S6 - Changes in mode occupancy between Mild Cognitive Impairment (MCI) and Dementia (DEM).** Barplots showing occupancy in MCI and DEM, highlighting differences in red for $p_{perm} < 0.05$, green for $p_{perm} < 0.05/K$, and blue for $p_{perm} < 2.39 \times 10^{-4}$.

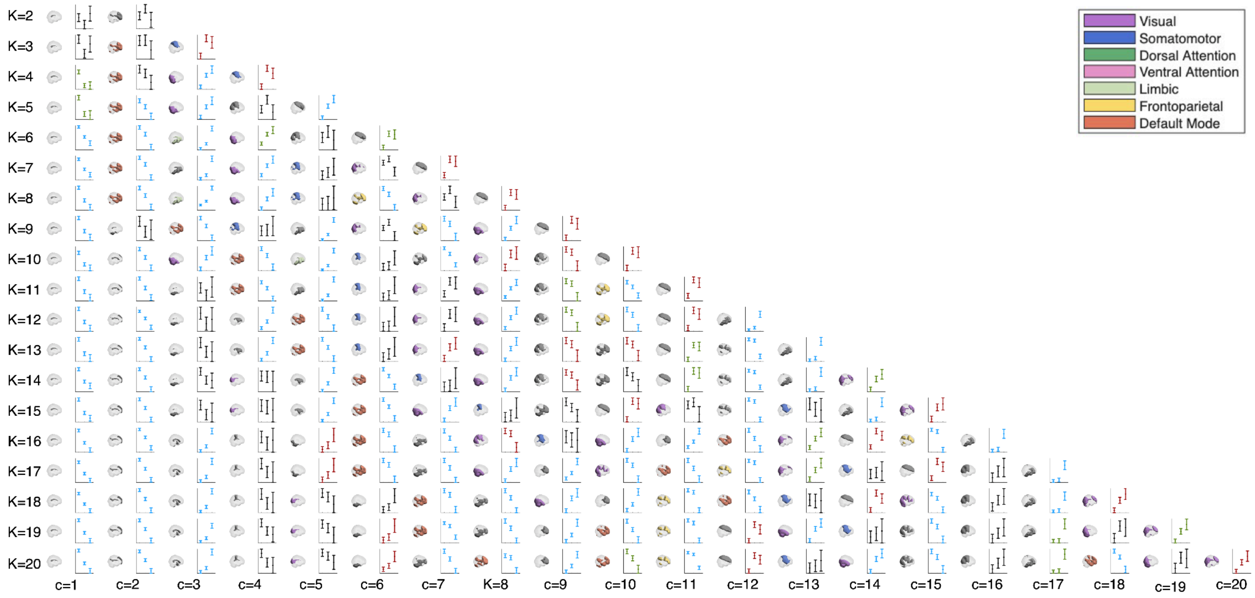


**Supplementary Figure S7 - Coupling modes detected with LEiDA across all 2177 fMRI scans, comparing their occurrence between CN, MCI and DEM.** a. Pyramid showing all the 209 phase coupling modes detected when applying k-means clustering to the entire sample of leading eigenvectors (from 2177 fMRI scans of 880 unique subjects), with the number of clusters K increasing from K = 2 (top) to K = 20 (bottom) clusters. Each mode is represented by the corresponding cluster centroid (c) shown in a transparent brain viewed from the side. Modes are colored according to the RSN defined by Yeo et al. (2011) with Dice coefficient >0.5, or in grey otherwise. Bar plots report the mean occupancy (± SEM) for all participants of each group, colored according to the most significant pairwise comparison between groups: red $p_{perm} < 0.05$, green $p_{perm} < 0.05/K$, blue for $p_{perm} < 2.39 \times 10\text{-}4$, or black when not significant ($p_{perm} > 0.05$).

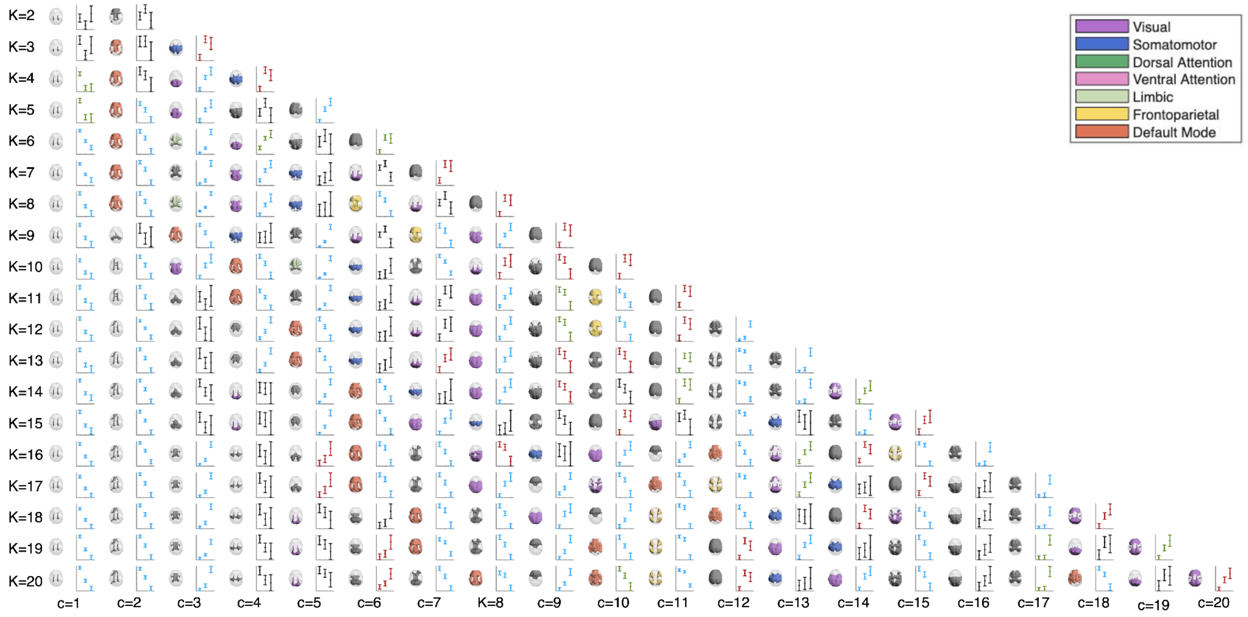


**Supplementary Figure S8 - Coupling modes detected with LEiDA across all 2177 fMRI scans, comparing their occurrence between CN, MCI and DEM.** Same as Supplementary Figure S7 but viewed from the top.

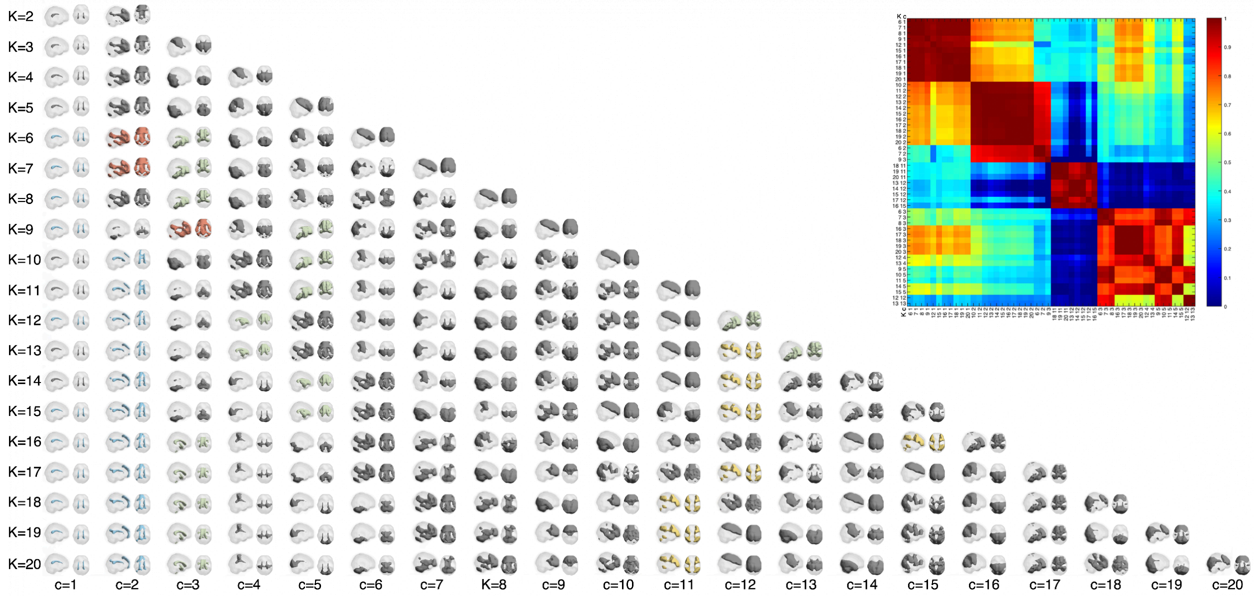


**Supplementary Figure S9 - Grouping into consistent Brain-Ventricle (BraVe) phase-coupling modes.** Pyramid of cluster centroids obtained for K = 2 to K = 20, where each row shows one clustering solution and each column (c = 1, …, K) one centroid within it. Each centroid is rendered on a transparent brain, viewed from the top and the side, highlighting voxels whose fMRI signals are dephased by more than 90° relative to the global signal direction. Clusters whose probability of occurrence differed significantly between any pair of conditions with $p < 0.05 / (209 \times 2 \times 3)$ and effect size > 0.35 were divided into groups with strong spatial overlap (Pearson's correlation >0.65): **BraVe mode I (blue, most significant for K=19, c=1)**, **BraVe mode II (orange, most significant for K=7, c=2)**, **BraVe mode III (yellow, most significant for K=20, c=11)**, and **BraVe mode IV (green, most significant for K=8, c=3)**; all remaining centroids are shown in gray. **Top-right:** Pearson's correlation matrix computed between every pair of the 50 cluster centroids that survived the significance and effect-size criteria in any pairwise comparison between conditions, highlighting the subdivision into 4 groups

**Supplementary Table S3. Permutation test statistics for BraVe mode occupancy group comparisons across three analysis samples.** p-values from permutation testing with 100,000 permutations. Differences surviving Bonferroni-corrected significance threshold ($p < 0.00417$, correcting for 3 pairwise comparisons × 4 BraVe modes) in bold font.

| **BraVe mode** | **Comparison** | **All scans (N=2177)** | | **First scan (N=880)** | | **Last scan (N=880)** | |
|---|---|---|---|---|---|---|---|
| | | p-value | Hedges' g | p-value | Hedges' g | p-value | Hedges' g |
| BraVe I | CN vs MCI | **2.38e-08** | **0.29** | 0.00582 | 0.21 | 0.00053 | 0.27 |
| BraVe I | CN vs DEM | **2.10e-08** | **0.46** | **2.65e-07** | **0.49** | **2.35e-07** | **0.57** |
| BraVe I | MCI vs DEM | 0.00421 | 0.20 | 0.00589 | 0.32 | 0.00738 | 0.29 |
| BraVe II | CN vs MCI | 0.00756 | 0.12 | 0.0442 | 0.15 | 0.0102 | 0.20 |
| BraVe II | CN vs DEM | **2.43e-08** | **0.37** | **0.000379** | **0.40** | **4.56e-07** | **0.39** |
| BraVe II | MCI vs DEM | **0.000189** | **0.25** | 0.0287 | 0.26 | 0.0514 | 0.20 |
| BraVe III | CN vs MCI | 0.324 | 0.05 | 0.340 | 0.07 | 0.0673 | 0.14 |
| BraVe III | CN vs DEM | **2.25e-08** | **0.42** | **1.93e-05** | **0.48** | **2.57e-07** | **0.45** |
| BraVe III | MCI vs DEM | **2.47e-08** | **0.38** | **0.000418** | **0.41** | **0.00181** | **0.34** |
| BraVe IV | CN vs MCI | **0.00130** | **0.15** | 0.434 | 0.06 | 0.102 | 0.13 |
| BraVe IV | CN vs DEM | **2.01e-08** | **0.63** | **0.000168** | **0.52** | **2.61e-07** | **0.66** |
| BraVe IV | MCI vs DEM | **2.89e-08** | **0.46** | **0.000368** | **0.43** | **2.66e-07** | **0.51** |

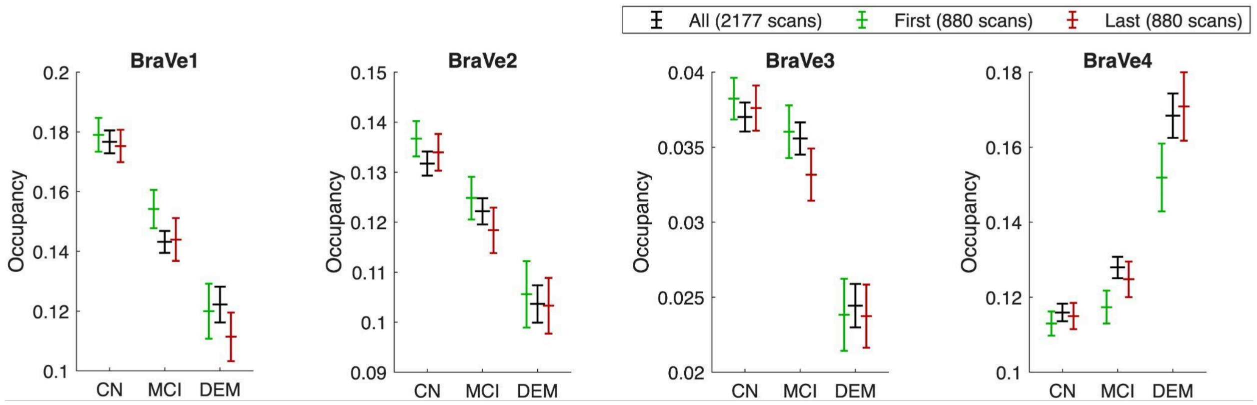


**Supplementary Figure S10. Replication of BraVe mode occupancy group differences in single-subject subsets.** Mean fractional occupancy (± SEM) of the four key BraVe modes across diagnostic groups (CN: cognitively normal; MCI: mild cognitive impairment; DEM: dementia) for three analysis samples: all available scans (black, N=2177), first scan per subject (green, N=880), and last scan per subject (red, N=880). The last scan was selected to ensure diagnostic labels reflect the most advanced clinical state captured during follow-up; the first scan was selected to test whether BraVe signatures are detectable prior to disease progression. Group differences were assessed using permutation testing (100,000 permutations) with Bonferroni correction for multiple comparisons. All four modes replicate the monotonic ordering across diagnostic groups in both single-subject subsets, with effect sizes and significance levels consistent with the full longitudinal sample (see Supplementary Table S3). Error bars represent SEM across subjects.

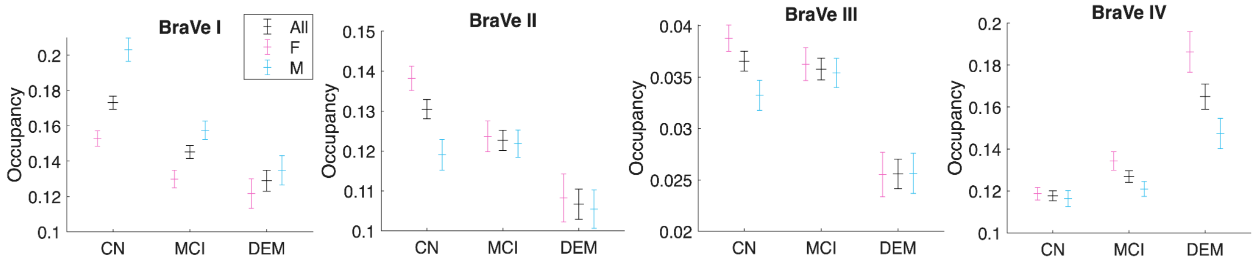


**Supplementary Figure S11 - Sex-stratified BraVe mode occupancy corrected for age and education.** Sex differences in BraVe mode occupancy after correction for age and education. Mean occupancy (± SEM) for all participants (black), females (pink), and males (cyan) across cognitive groups. Occupancy values were harmonized across sites using ComBat with age, sex, and education as protected covariates, then corrected for age and education via linear regression. Asterisks indicate effects surviving Bonferroni correction across four modes ($p < 0.0125$). See Supplementary Table S4 for full statistics. Occupancy values were additionally corrected for age and education via linear regression, with the grand mean added back to preserve the original occupancy scale, to ensure that the plotted group means and standard errors reflect occupancy values independent of both confounds.

**Supplementary Table S4. Two-way ANCOVA results for the effect of sex on BraVe mode occupancy, with age and years of education included as continuous covariates.** ✓ indicates $p < 0.0125$ (Bonferroni correction across four modes). Education differed significantly between sexes (females 15.9 vs males 16.6 years, $p = 1.2 \times 10^{-11}$).

| Effect | **Mode I** | | **Mode II** | | **Mode III** | | **Mode IV** | |
|---|---|---|---|---|---|---|---|---|
| **Condition** | 2.87E-16 | ✓ | 4.88E-05 | ✓ | 2.67E-07 | ✓ | 1.56E-19 | ✓ |
| **Sex** | 5.33E-09 | ✓ | 3.66E-02 | | 1.78E-01 | | 2.05E-06 | ✓ |
| **Age** | 2.84E-05 | ✓ | 1.71E-10 | ✓ | 5.02E-07 | ✓ | 1.70E-04 | ✓ |
| **Education** | 3.67E-04 | ✓ | 5.15E-01 | | 1.99E-01 | | 1.18E-01 | |
| **Condition x Sex** | 2.12E-02 | | 3.26E-02 | | 1.65E-01 | | 2.08E-03 | ✓ |

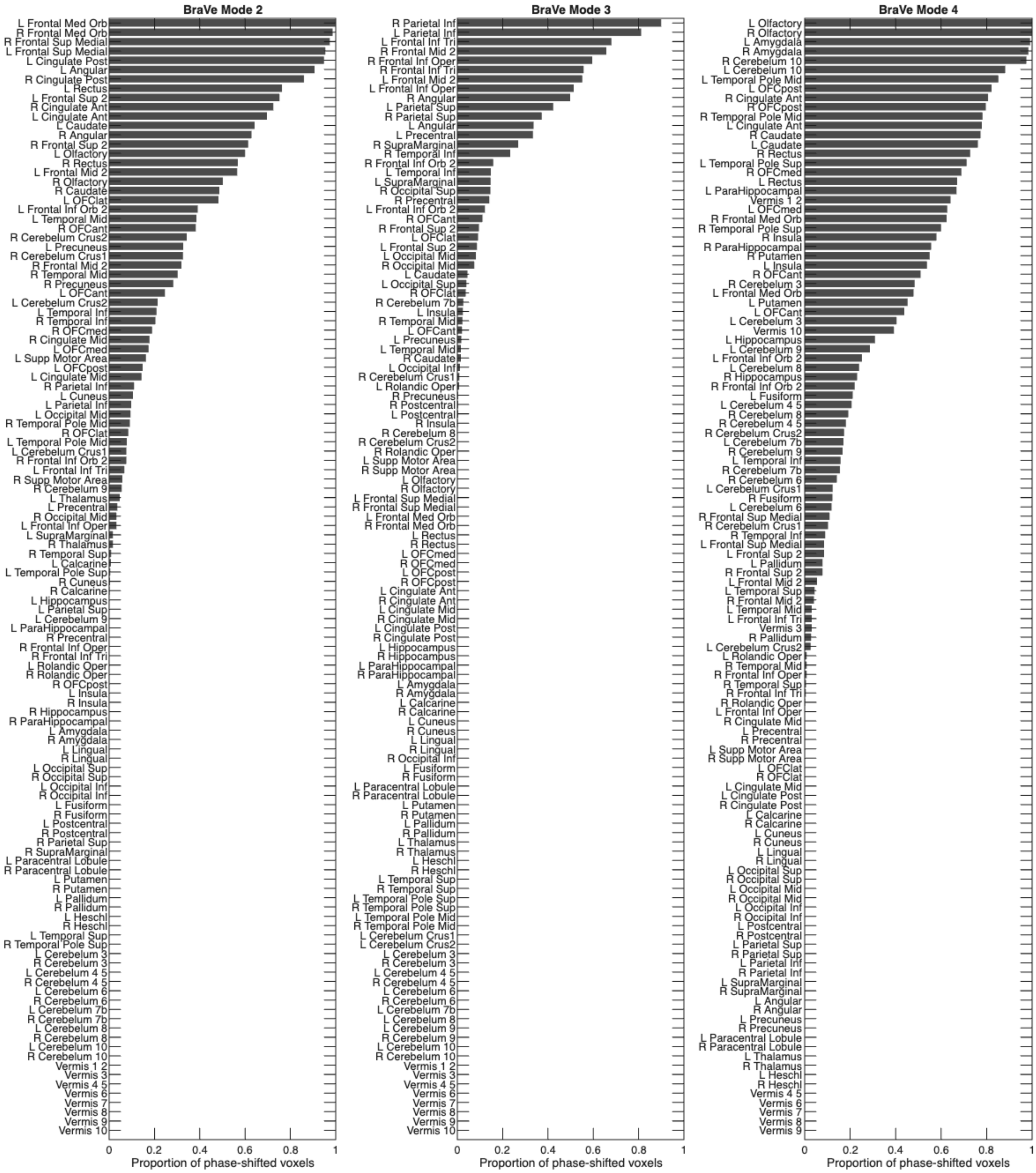


**Supplementary Figure S12 -** List of the regions from the Automated Anatomical Labeling (AAL, version 2) template, indicating the number of voxels whose signals shift by more than pi/2 from the rest of the brain in each of the 3 complex Brain-Ventricle Phase coupling modes detected: BraVe mode II (**a**), BraVe mode III (**b**), and, BraVe mode IV (**c**). The list of brain areas and their full names is provided below.

#### List of brain areas in the AAL2 template

1. Precentral Gyrus, Left (L Precentral)
2. Precentral Gyrus, Right (R Precentral)
3. Superior Frontal Gyrus 2, Left (L Frontal Sup 2)
4. Superior Frontal Gyrus 2, Right (R Frontal Sup 2)
5. Middle Frontal Gyrus 2, Left (L Frontal Mid 2)
6. Middle Frontal Gyrus 2, Right (R Frontal Mid 2)
7. Inferior Frontal Gyrus, Opercular Part, Left (L Frontal Inf Oper)
8. Inferior Frontal Gyrus, Opercular Part, Right (R Frontal Inf Oper)
9. Inferior Frontal Gyrus, Triangular Part, Left (L Frontal Inf Tri)
10. Inferior Frontal Gyrus, Triangular Part, Right (R Frontal Inf Tri)
11. Inferior Frontal Gyrus, Orbital Part 2, Left (L Frontal Inf Orb 2)
12. Inferior Frontal Gyrus, Orbital Part 2, Right (R Frontal Inf Orb 2)
13. Rolandic Operculum, Left (L Rolandic Oper)
14. Rolandic Operculum, Right (R Rolandic Oper)
15. Supplementary Motor Area, Left (L Supp Motor Area)
16. Supplementary Motor Area, Right (R Supp Motor Area)
17. Olfactory Cortex, Left (L Olfactory)
18. Olfactory Cortex, Right (R Olfactory)
19. Superior Frontal Gyrus, Medial Part, Left (L Frontal Sup Medial)
20. Superior Frontal Gyrus, Medial Part, Right (R Frontal Sup Medial)
21. Medial Orbital Frontal Cortex, Left (L Frontal Med Orb)
22. Medial Orbital Frontal Cortex, Right (R Frontal Med Orb)
23. Gyrus Rectus, Left (L Rectus)
24. Gyrus Rectus, Right (R Rectus)
25. Medial Orbital Frontal Cortex, Left (L OFCmed)
26. Medial Orbital Frontal Cortex, Right (R OFCmed)
27. Anterior Orbital Frontal Cortex, Left (L OFCant)
28. Anterior Orbital Frontal Cortex, Right (R OFCant)
29. Posterior Orbital Frontal Cortex, Left (L OFCpost)
30. Posterior Orbital Frontal Cortex, Right (R OFCpost)
31. Lateral Orbital Frontal Cortex, Left (L OFClat)
32. Lateral Orbital Frontal Cortex, Right (R OFClat)
33. Insula, Left (L Insula)
34. Insula, Right (R Insula)
35. Anterior Cingulate Gyrus, Left (L Cingulate Ant)
36. Anterior Cingulate Gyrus, Right (R Cingulate Ant)
37. Middle Cingulate Gyrus, Left (L Cingulate Mid)
38. Middle Cingulate Gyrus, Right (R Cingulate Mid)
39. Posterior Cingulate Gyrus, Left (L Cingulate Post)
40. Posterior Cingulate Gyrus, Right (R Cingulate Post)
41. Hippocampus, Left (L Hippocampus)
42. Hippocampus, Right (R Hippocampus)
43. Parahippocampal Gyrus, Left (L ParaHippocampal)
44. Parahippocampal Gyrus, Right (R ParaHippocampal)
45. Amygdala, Left (L Amygdala)
46. Amygdala, Right (R Amygdala)
47. Calcarine Fissure and Surrounding Cortex, Left (L Calcarine)
48. Calcarine Fissure and Surrounding Cortex, Right (R Calcarine)
49. Cuneus, Left (L Cuneus)
50. Cuneus, Right (R Cuneus)
51. Lingual Gyrus, Left (L Lingual)
52. Lingual Gyrus, Right (R Lingual)
53. Superior Occipital Gyrus, Left (L Occipital Sup)
54. Superior Occipital Gyrus, Right (R Occipital Sup)
55. Middle Occipital Gyrus, Left (L Occipital Mid)
56. Middle Occipital Gyrus, Right (R Occipital Mid)
57. Inferior Occipital Gyrus, Left (L Occipital Inf)
58. Inferior Occipital Gyrus, Right (R Occipital Inf)
59. Fusiform Gyrus, Left (L Fusiform)
60. Fusiform Gyrus, Right (R Fusiform)
61. Postcentral Gyrus, Left (L Postcentral)
62. Postcentral Gyrus, Right (R Postcentral)
63. Superior Parietal Gyrus, Left (L Parietal Sup)
64. Superior Parietal Gyrus, Right (R Parietal Sup)
65. Inferior Parietal Lobule, Left (L Parietal Inf)
66. Inferior Parietal Lobule, Right (R Parietal Inf)
67. Supramarginal Gyrus, Left (L SupraMarginal)
68. Supramarginal Gyrus, Right (R SupraMarginal)
69. Angular Gyrus, Left (L Angular)
70. Angular Gyrus, Right (R Angular)
71. Precuneus, Left (L Precuneus)
72. Precuneus, Right (R Precuneus)
73. Paracentral Lobule, Left (L Paracentral Lobule)
74. Paracentral Lobule, Right (R Paracentral Lobule)
75. Caudate Nucleus, Left (L Caudate)
76. Caudate Nucleus, Right (R Caudate)
77. Putamen, Left (L Putamen)
78. Putamen, Right (R Putamen)
79. Globus Pallidus, Left (L Pallidum)
80. Globus Pallidus, Right (R Pallidum)
81. Thalamus, Left (L Thalamus)
82. Thalamus, Right (R Thalamus)
83. Heschl's Gyrus (Transverse Temporal Gyrus), Left (L Heschl)
84. Heschl's Gyrus (Transverse Temporal Gyrus), Right (R Heschl)
85. Superior Temporal Gyrus, Left (L Temporal Sup)
86. Superior Temporal Gyrus, Right (R Temporal Sup)
87. Superior Temporal Pole, Left (L Temporal Pole Sup)
88. Superior Temporal Pole, Right (R Temporal Pole Sup)
89. Middle Temporal Gyrus, Left (L Temporal Mid)
90. Middle Temporal Gyrus, Right (R Temporal Mid)
91. Middle Temporal Pole, Left (L Temporal Pole Mid)
92. Middle Temporal Pole, Right (R Temporal Pole Mid)
93. Inferior Temporal Gyrus, Left (L Temporal Inf)
94. Inferior Temporal Gyrus, Right (R Temporal Inf)
95. Cerebellum Crus I, Left (L Cerebelum Crus1)
96. Cerebellum Crus I, Right (R Cerebelum Crus1)
97. Cerebellum Crus II, Left (L Cerebelum Crus2)
98. Cerebellum Crus II, Right (R Cerebelum Crus2)
99. Cerebellum Lobule III, Left (L Cerebelum 3)
100. Cerebellum Lobule III, Right (R Cerebelum 3)
101. Cerebellum Lobules IV-V, Left (L Cerebelum 4 5)
102. Cerebellum Lobules IV-V, Right (R Cerebelum 4 5)
103. Cerebellum Lobule VI, Left (L Cerebelum 6)
104. Cerebellum Lobule VI, Right (R Cerebelum 6)
105. Cerebellum Lobule VIIb, Left (L Cerebelum 7b)
106. Cerebellum Lobule VIIb, Right (R Cerebelum 7b)
107. Cerebellum Lobule VIII, Left (L Cerebelum 8)
108. Cerebellum Lobule VIII, Right (R Cerebelum 8)
109. Cerebellum Lobule IX, Left (L Cerebelum 9)
110. Cerebellum Lobule IX, Right (R Cerebelum 9)
111. Cerebellum Lobule X, Left (L Cerebelum 10)
112. Cerebellum Lobule X, Right (R Cerebelum 10)
113. Cerebellar Vermis Segments I/II (Vermis 1 2)
114. Cerebellar Vermis Segment III (Vermis 3)
115. Cerebellar Vermis Segments IV/V (Vermis 4 5)
116. Cerebellar Vermis Segment VI (Vermis 6)
117. Cerebellar Vermis Segment VII (Vermis 7)
118. Cerebellar Vermis Segment VIII (Vermis 8)
119. Cerebellar Vermis Segment IX (Vermis 9)
120. Cerebellar Vermis Segment X (Verm

**Supplementary Table S5 - Pearson's correlation between Cognitive Scores and Biomarkers vs Mode Occupancies.** Raw p-values are reported, Bonferroni correction threshold for true positives is $3.47\times10^{-4}$, correcting for 36 variables × 4 modes = 144 tests:

| | BraVe Mode I | | BraVe Mode II | | BraVe Mode III | | BraVe Mode IV | |
|---|---|---|---|---|---|---|---|---|
| **Variable** | **Pearson r** | **p-value** | **Pearson r** | **p-value** | **Pearson r** | **p-value** | **Pearson r** | **p-value** |
| **APOE4 (N=2041)** | -0,056 | 1,44E-02 | 0,011 | 6,29E-01 | -0,012 | 5,96E-01 | 0,110 | 1,3E-06 |
| **FDG (N=444)** | 0,079 | 1,08E-01 | 0,045 | 3,59E-01 | 0,200 | 3,65E-05 | -0,067 | 1,73E-01 |
| **AV45 (N=911)** | -0,071 | 3,95E-02 | 0,001 | 9,76E-01 | -0,045 | 1,95E-01 | 0,097 | 4,84E-03 |
| **FBB (N=304)** | -0,134 | 2,52E-02 | -0,012 | 8,38E-01 | -0,018 | 7,62E-01 | 0,026 | 6,68E-01 |
| **ABETA (N=207)** | 0,051 | 4,65E-01 | -0,016 | 8,21E-01 | 0,070 | 3,17E-01 | -0,031 | 6,59E-01 |
| **TAU (N=263)** | -0,050 | 4,27E-01 | -0,077 | 2,2E-01 | -0,032 | 6,08E-01 | 0,076 | 2,26E-01 |
| **PTAU (N=261)** | -0,032 | 6,08E-01 | -0,083 | 1,89E-01 | -0,039 | 5,32E-01 | 0,060 | 3,36E-01 |
| **MMSE (N=2012)** | 0,116 | 5,23E-07 | 0,086 | 1,78E-04 | 0,122 | 1,24E-07 | -0,162 | 1,97E-12 |
| **MOCA (N=1960)** | 0,154 | 3,61E-11 | 0,075 | 1,34E-03 | 0,136 | 5,74E-09 | -0,162 | 3,2E-12 |
| **LDELTOTAL (N=1862)** | 0,095 | 7,22E-05 | 0,088 | 2,69E-04 | 0,138 | 7,63E-09 | -0,125 | 1,82E-07 |
| **mPACCdigit (N=2019)** | 0,127 | 3,42E-08 | 0,100 | 1,35E-05 | 0,142 | 5,62E-10 | -0,175 | 1,87E-14 |
| **mPACCtrailsB (N=2019)** | 0,145 | 2,64E-10 | 0,103 | 7,28E-06 | 0,146 | 2,14E-10 | -0,186 | 3,92E-16 |
| **RAVLT_immediate (N=1999)** | 0,102 | 1,07E-05 | 0,101 | 1,31E-05 | 0,119 | 2,85E-07 | -0,141 | 1,06E-09 |
| **RAVLT_learning (N=1999)** | 0,096 | 3,1E-05 | 0,095 | 4,21E-05 | 0,094 | 4,41E-05 | -0,126 | 4,33E-08 |
| **RAVLT_forgetting (N=1996)** | -0,003 | 8,8E-01 | 0,011 | 6,31E-01 | -0,039 | 9,03E-02 | 0,041 | 7,68E-02 |
| **RAVLT_perc_forgetting (N=1995)** | -0,060 | 9,14E-03 | -0,012 | 6,03E-01 | -0,075 | 1,17E-03 | 0,103 | 8,94E-06 |
| **CDRSB (N=2002)** | -0,141 | 9,04E-10 | -0,115 | 5,85E-07 | -0,116 | 5,52E-07 | 0,202 | 1,13E-18 |
| **ADAS11 (N=2006)** | -0,131 | 1,17E-08 | -0,109 | 2,17E-06 | -0,135 | 5,12E-09 | 0,200 | 2,78E-18 |
| **ADAS13 (N=1998)** | -0,140 | 1,09E-09 | -0,111 | 1,51E-06 | -0,136 | 3,94E-09 | 0,196 | 1,16E-17 |
| **ADASQ4 (N=2015)** | -0,119 | 2,14E-07 | -0,082 | 3,58E-04 | -0,114 | 8,07E-07 | 0,144 | 3,16E-10 |
| **TRABSCOR (N=1949)** | -0,159 | 1,05E-11 | -0,061 | 9,77E-03 | -0,090 | 1,17E-04 | 0,164 | 2,26E-12 |
| **EcogPtMem (N=1999)** | -0,111 | 1,42E-06 | -0,070 | 2,54E-03 | -0,031 | 1,78E-01 | 0,063 | 6,26E-03 |
| **EcogPtLang (N=1995)** | -0,110 | 2,18E-06 | -0,039 | 9,63E-02 | -0,014 | 5,56E-01 | 0,070 | 2,59E-03 |
| **EcogPtVisspat (N=1967)** | -0,136 | 5,58E-09 | -0,055 | 1,94E-02 | -0,056 | 1,7E-02 | 0,087 | 1,77E-04 |
| **EcogPtPlan (N=1997)** | -0,124 | 7,79E-08 | -0,066 | 4,22E-03 | -0,060 | 1,01E-02 | 0,063 | 6,78E-03 |
| **EcogPtOrgan (N=1954)** | -0,123 | 1,49E-07 | -0,109 | 3,14E-06 | -0,065 | 5,68E-03 | 0,034 | 1,44E-01 |
| **EcogPtDivatt (N=1984)** | -0,099 | 2,07E-05 | -0,050 | 3,25E-02 | -0,005 | 8,18E-01 | 0,055 | 1,76E-02 |
| **EcogPtTotal (N=1999)** | -0,136 | 4,1E-09 | -0,073 | 1,59E-03 | -0,037 | 1,09E-01 | 0,074 | 1,33E-03 |
| **EcogSPMem (N=1974)** | -0,121 | 2,2E-07 | -0,101 | 1,45E-05 | -0,117 | 4,85E-07 | 0,155 | 2,37E-11 |
| **EcogSPLang (N=1973)** | -0,113 | 1,33E-06 | -0,079 | 7,44E-04 | -0,092 | 8,07E-05 | 0,140 | 1,49E-09 |
| **EcogSPVisspat (N=1926)** | -0,124 | 1,54E-07 | -0,090 | 1,35E-04 | -0,108 | 4,87E-06 | 0,152 | 9,09E-11 |
| **EcogSPPlan (N=1958)** | -0,118 | 4,38E-07 | -0,088 | 1,7E-04 | -0,103 | 9,66E-06 | 0,143 | 8,44E-10 |
| **EcogSPOrgan (N=1906)** | -0,140 | 3,62E-09 | -0,121 | 2,91E-07 | -0,114 | 1,47E-06 | 0,157 | 2,69E-11 |
| **EcogSPDivatt (N=1929)** | -0,104 | 9,9E-06 | -0,089 | 1,5E-04 | -0,094 | 6,16E-05 | 0,139 | 2,97E-09 |
| **EcogSPTotal (N=1973)** | -0,126 | 5,42E-08 | -0,105 | 7,22E-06 | -0,115 | 7,52E-07 | 0,160 | 6,14E-12 |

### Supplementary References

1. Cabral J, Vidaurre D, Marques P, et al. (2017). Cognitive performance in healthy older adults relates to spontaneous switching between states of functional connectivity during rest. Scientific Reports, 7(1), 1-9.
2. Fortin J-P, Parker D, Tunc B, et al. (2017). Harmonization of multi-site diffusion tensor imaging data. NeuroImage, 161, 149-170.
3. Fortin J-P, Cullen N, Sheline YI, et al. (2018). Harmonization of cortical thickness measurements across scanners and sites. NeuroImage, 167, 104-120.
4. Gorgolewski KJ, Auer T, Calhoun VD, et al. (2016). The brain imaging data structure, a format for organizing and describing outputs of neuroimaging experiments. Scientific Data, 3, 160044.
5. Johnson WE, Li C, Rabinovic A (2007). Adjusting batch effects in microarray expression data using empirical Bayes methods. Biostatistics, 8(1), 118-127.
6. Mueller SG, Weiner MW, Thal LJ, et al. (2005). The Alzheimer's Disease Neuroimaging Initiative. Neuroimaging Clinics of North America, 15(4), 869-877.
